# Impact of co-occurrent assortative mating and vertical cultural transmission on measures of genetic associations

**DOI:** 10.1101/2023.04.08.536101

**Authors:** Anthony F. Herzig, Camille Noûs, Aude Saint Pierre, Hervé Perdry

## Abstract

Assortative mating for a given phenotype is the phenomenon by which mates select each other based on their phenotypic similarity. Other phenomena can create positive correlation between the parents’ and the offspring’s environment: vertical cultural transmission, or dynastic effects. When these phenomena occur together, they induce a gene-environment correlation at the population scale. It will impact genetic measures of associations such as SNP effect size and SNP-heritability.

In this paper, we provide a complete mathematical modelling of both assortative mating and vertical cultural transmission in the classical framework of the polygenic additive model. We establish for the first time the theoretical evolution and equilibrium values of all quantities of interest involved. We then derive its consequences on typical genetic epidemiology analyses; including both population and family-based study designs. We show the consequences on heritability estimation, Genome-Wide Association Studies (GWAS), and the variance explained by polygenic scores. We validate our calculations through simple forward-time simulations.

## 1 Introduction

Assortative mating (AM) is the process of non-random mate choice that results in a positive correlation of phenotypic values and was first described by RA Fisher in his seminal presentation of the polygenic model in 1919 (Fisher, 1919). The theoretical consequences of AM on the distribution of genetic variants in a population have been explored in detail, notably by Crow and Felsenstein (Crow & Felsenstein, 1968), Wright (Wright, 1921) and Nagylaki (Nagylaki, 1978). Importantly, it has been shown that AM results in long distance correlations between genetic variants that have an effect on the phenotype (under a polygenic model) and that this will result in an increase in both phenotypic and breeding (additive genetic)-value variances.

Correlated phenotypes between mates have been observed directly in human populations. A most simple example is the often-studied correlation between standing height of spouses (Pearson & Lee, 1903; Stulp et al., 2017). Correlated phenotypes have also been regularly estimated for a range of social and behavioural traits (Buss, 1985) including education level (Hugh-Jones et al., 2016; Mare, 1991) as well as for multifactorial disease (Hippisley-Cox et al., 2002; Yamamoto et al., 2023) and psychiatric disorders (de Jong et al., 2018; Jefsen et al., 2022; Nordsletten et al., 2016).

In the age of widely available genetic data and increasingly huge cross-sectional epidemiological studies including genetic data (All of Us Research Program Investigators et al., 2019; Bycroft et al., 2018; Nagai et al., 2017), it has become readily possible to search for the signs of AM on genetic variation in a population (Border, Athanasiadis, et al., 2022; Yamamoto et al., 2023; Yengo et al., 2018), validating some predictions made by early population geneticists almost 100 years later. Such findings have indicated that AM is likely prevalent and capable of distorting the inference of prevalent genetic epidemiology study design such as genome-wide association studies (GWAS) (Howe et al., 2021, 2022), heritability analysis (Border, O’Rourke, et al., 2022), polygenic score prediction (Torvik et al., 2022; Yengo et al., 2018) and Mendelian randomisation (Brumpton et al., 2020; Davies et al., 2019; Hartwig et al., 2018; Howe et al., 2019). Indeed, assortative mating has been recently described as one of the principal confounders of genetic association studies (Morris et al., 2020; Veller & Coop, 2023).

A further phenomenon that impacts studies in genetic epidemiological studies is that of vertical cultural transmission (VCT), in which environment is transmitted from parents to offspring. This can be simply modelled as a (positive) correlation between parents and offspring environments. Such VCT can intuitively produce correlations between phenotypes that can closely mimic the correlation induced by an additive genetic component (Cavalli-Sforza & Feldman, 1973). Indeed, the importance of considering the non-independence of non-genetic factors between closely related individuals has long been recognized and is a motivation for the introduction of twin studies in estimating heritability (Falconer, 1960).

A similar notion to VCT is that of a dynasty effect (DE) as described in (Brumpton et al., 2020) and in (Morris et al., 2020). VCT and DE respectively correspond to the influence of parental environments or parental phenotypes on offspring environment; both leading to a positive correlation between the non-genetic contributions to phenotypic variance between parents and offspring. Such effects were characterised by the term ‘the Nature of Nurture’ in (Kong et al., 2018) capturing the idea that parental genotypes (or even genotypes of more distant relatives) can affect the phenotype of the offspring though paths other than the actual sharing of alleles. Various methods have been put forward to control for such structure when estimating heritability (A. I. Young et al., 2018; Zaitlen et al., 2013) or GWAS (Howe et al., 2022; A. I. Young et al., 2022) based on the idea of controlling for parental genotypes in order to try to capture the direct substitution effects of genetic factors.

Whilst both AM and VCT have therefore been widely addressed in terms of their impact of population based genetic epidemiology studies, there is no reason to suppose that AM and VCT may not co-occur. A general model that takes into account the combined consequences of AM and VCT was set out by Rice, Cloninger and Reich in three linked papers in 1978-79 (Cloninger et al., 1979a, 1979b; Rice et al., 1978). Rather than explicitly describing polygenic inheritance as in Crow and Felsenstein, 1968, path analysis is used by Rice, Cloninger and Reich with the simple assumption of the path coefficient between the genetic component of a parent and the offspring being ½ (Rice et al., 1978). Crucially, Rice, Cloninger and Reich describe the correlation between genetic and cultural components within an individual; *w* in their notation, *ρ* in ours. The possible presence of this correlation was first explored (to the best of our knowledge) in path analyses in (Rao et al., 1976). Gene-environment correlations involving models of complex gene-culture transmission and assortative mating are also presented in (Feldman & Cavalli-Sforza, 1977) and (Cavalli-Sforza & Feldman, 1978). Rice, Cloninger and Reich describe such gene-environment correlations as being either a primary correlation (a characteristic of the population) or one that can arise and be derived in the presence of AM and VCT (Cloninger et al., 1979a). This correlation was explored in real data by Rice, Cloninger and Reich for IQ scores and social-economic status data in the USA (Rice et al., 1980); estimating a value for *w* between 0.137 and 0.191. This correlation between genetics and cultural effects within individuals subsequently gained some traction in the field of behavioural genetics; for a few examples it is included in the model in (Carey & Rice, 1983) where it is explicitly described as being induced by AM and VCT; as well as in many subsequent works: for example in (Vogler & Fulker, 1983), (Martin et al., 1986), (Truett et al., 1994), and more recently in (Keller et al., 2009), (Vinkhuyzen et al., 2012), and (Balbona et al., 2022). It may be the case that such complicated models involving AM and VCT have found their niche in the analysis of highly complex social traits (e.g. intelligence and personality traits) where complex familial data (twin, siblings, adoption studies etc.) is often leveraged. The potential of an important impact on co-occurrent AM and VCT on twin studies in general was recently discussed in (Larrègue et al., 2023) for behavioural traits.

More broadly, in genetic epidemiological studies based on biobanks, any such complications are generally ignored and environmental/cultural factors are assumed to act independently from genetics. We therefore wished to revisit the question of co-occurring AM and VCT and explore the implication of the genetic-cultural correlation in modern study designs in genetic epidemiology. To this end, we derive here (and confirm through simulation) the evolution and equilibrium equations for the correlation structure between genetic and environmental values in a trio in the presence of both AM and VCT. Unlike previous work, we achieve this with an explicit model of polygenic inheritance including derivations of the eventual correlations that occur between all pairs of causal genetic variants in a population. Indeed, this allows us to calculate evolution and equilibrium values that to our knowledge have never been derived even in the simpler model involving only AM. Our key result is to show that the combination of AM and VCT leads to positive correlation between genetic and environmental values at the population level and that this has significant impact on subsequent study designs. Specifically, this leads to inflated SNP-heritability and SNP-effect estimates, and hence also on polygenic score (PGS)-based methods, due to the fact that estimated genetic effects will also capture non-genetic effects. We also investigate the impact on study designs based on analysing large-scale family data. Simple forward-time simulations are presented to validate our calculations and demonstrate the impact of co-occurring AM and VCT on the measures of association.

## 2 Methods

### 2.1 The polygenic additive model with assortative mating and vertical cultural transmission (AMVCT)

#### 2.1.1 Model without homogamy

We consider Fisher’s infinitesimal additive model, where a phenotype is determined by *P* =*A* +*E* where the genetic value *A* is the sum of the effects of a large number *N* of genetic loci and *E* is the environmental effect. We consider two mates indexed by 1 and 2, and an offspring index by 3.

Denote var(*A*) =*a*^2^, var(*E*) =*e*^2^ and var(*P*) =*σ*^2^ =*a*^2^+ *e*^2^. If the population is panmictic, and assuming *E*_1_, *E*_2_ and *E*_3_ are independent, the correlation between the genetic values of the parent and the offspring is classically 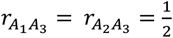(Figure 1, panel a), leading to phenotypic correlations 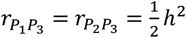 where 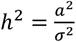, classically referred to as the (narrow-sense) heritability.

**Figure 1.**
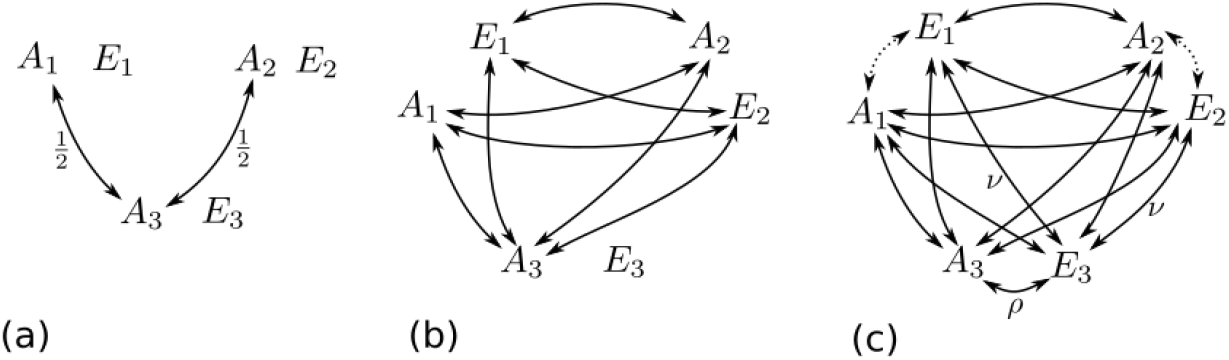
The correlations between genetic values and environments of parents (indexed by 1, 2) and offspring (indexed by 3), in models of increasing complexity. Panel (a) corresponds to the simple model of a panmictic population; (b) corresponds to a model of homogamy, in which the positive phenotypic correlations between the mates induces positive correlations between *A*_1_ (respectively *E*_1_) and both *A*_2_ and *E*_2_, inducing correlations between *A*_3_ and all the four parental variables, but in which *E*_3_ is independent of all other variables; and (c) corresponds to the introduction of an additional parameter *v* for the correlation between the parental environment *E*_1_(respectively *E*_2_) and the offspring environment *E*_3_, inducing positive correlations between all pairs of variables, except possibly *A*_1_and *E*_1_ (respectively *A*_2_ and *E*_2_), if this is the first generation of random mating.

#### 2.1.2 Introducing homogamy

We consider a panmictic population in which homogamy appears at a generation indexed by *t* =0. Denote by 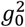 the gametic variance in the absence of homogamy and the heritability in the absence of homogamy as 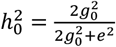. In the presence of homogamy, we denote 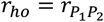 the correlation between the mates’ phenotypes. The presence of this correlation induces a correlation 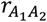 between *A*_1_and *A*_2_, so the genetic values of the gametes emitted by the parents have a positive correlation *r*_*ga*_. This gametic correlation will, across generations, induce a small gametic disequilibrium between the causal loci, which will cause an increase in the gametic genetic variance 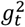 (and of the individual genetic variance 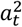), until an equilibrium is reached. At the same time, there is a correlation between *A*_1_and both *A*_2_ and *E*_2_, and also between *E*_1_and both *A*_2_ and *E*_2_. As the offspring genetic value *A*_3_ is correlated to *A*_1_ and *A*_2_, which are respectively correlated to *E*_2_ and *E*_1_, this implies the presence of a correlation between *A*_3_ and both *E*_1_ and *E*_2_ (Figure 1, panel b). The values of all these correlations at equilibrium will be computed (see below in 3.1.1, 3.1.2 and Supplementary Material sections 1-3).

#### 2.1.3 Introducing vertical cultural transmission

We consider the same model as above, but with yet another parameter *v* for the correlation between the parents’ environment values *E*_1_, *E*_2_, and the offspring environment *E*_3_ (Figure 1, panel c). As both *E*_1_and *E*_2_ are correlated to *A*_3_, this induces a correlation *ρ* between *A*_3_ and *E*_3_. Again, we will compute the se correlation in the Results and Supplementary Material sections 1-4. The value of *ρ* will increase generation after generation, until reaching an equilibrium point.

### 2.2 Forward-time Simulations

We performed simple simulations in R to demonstrate the validity of our evolution and equilibrium equations (simulation scripts are provided, see Data Availability). We simulated populations of constant size *M* =25,000 and 100,000 over subsequent generations with co-occurring AM and VCT. This involved simulating the population at generation 0, with a quantitative phenotype with variance *σ*^2^ =1 determined by *N* =1,000 unlinked causal SNPs and an architecture of 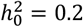 (hence, 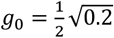 and 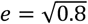). The gametic alleles were drawn with minor-allele frequencies of 0.5 and we applied equal allelic substitution effects for all causal SNPs.

AM was simulated by forming *M*/2 pairs in each generation, this was achieved by simulating a proxy variable for *M*/2 pairs from a bivariate normal distribution with a correlation of *r*_*ho*_, and then using the ranks across all *M* observations to form pairs with the simulated phenotype data. Each pair would produce two offspring to keep the generation size stable. Each causal SNP was transmitted randomly and independently of the others, i.e., with no linkage. VCT was simulated by drawing environmental components for each offspring (*E*_3_) using the parameter *v* to ensure the correct covariance structure between *E*_1_, *E*_2_ and *E*_3_ in each trio.

## 3 Results

### 3.1 Evolution and Equilibrium equations

#### 3.1.1 Correlations under AM

We computed the values of all correlations between components, both genetic and environmental, in a trio at equilibrium (see Supplementary Material sections 1-3) using the same approach as (Nagylaki, 1978). The non-zero correlations between the genetic and environmental values of the family members are summarized in table 1, where *a, e* and *σ* are the standard deviations of *A, E* and *P* at equilibrium. The gametic correlation verifies 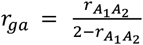, that is 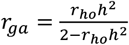. The gametic disequilibrium between each pair of the *N* causal loci corresponds to a correlation of 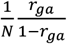; the cumulated effects of these intra gametic correlation leads to a gametic variance 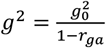. In turn, this leads to a genetic variance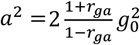. Additionally, we give in the Supplementary Material (sections 4 and 5; in particular section 4.5) equations for the evolution and equilibrium values of all these quantities across generations. Expressions for the equilibrium values for *r*_*ga*_ and *a*^2^ can be found that rely only on parameters of the initial population (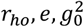); they are as follows:

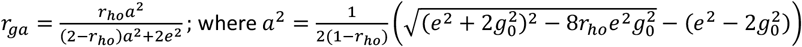

**Table 1.** Correlations between the genetic and the environmental effects in presence of assortative mating, without a shared parent-offspring environment. Individuals 1 and 2 play symmetric roles, so we have 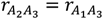 and 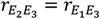.

| | $A_2$ | $E_2$ | $A_3$ |
| --- | --- | --- | --- |
| $A_1$ | $r_{ho}h^2$ | $r_{ho}\frac{ae}{\sigma^2}$ | $\frac{1}{2}(1 + r_{ho}h^2)$ |
| $E_1$ | $r_{ho}\frac{ae}{\sigma^2}$ | $r_{ho}(1 - h^2)$ | $\frac{1}{2}\left(1 + r_{ho}\frac{ae}{\sigma^2}\right)$ |

#### 3.1.2 Correlations arising from vertical cultural transmission

We give in the Supplementary Material evolution equations for all the quantities of interest for a model including VCT. Though there is no closed form for the equilibrium value of *ρ*, we show in Supplementary Material (section 4.4) that it can be computed by solving numerically a system of equations involving *ρ, v, e* (which does not change over time), and 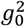. Considering that 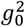 and *e* as fixed quantities, one gets that *ρ* is an increasing function of *v* with *ρ* =0 when *v* =0.

The gametic variance still verifies 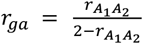 (with 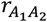 as in table 2 this time), and *g*^2^ and *a*^2^ depends on *r*_*ga*_ in the same way as before, thus in particular 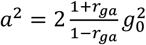. Table 2 summarizes the correlations between the genetic and environmental values of the family members, relying on the value of *ρ*.

**Table 2.**
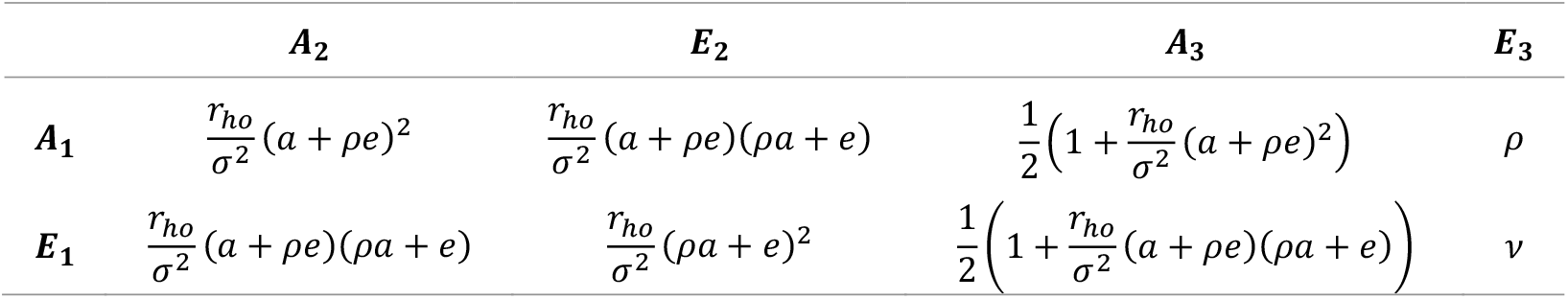
Correlations between the genetic and the environmental effects in presence of assortative mating, without shared parent-offspring environment. Individuals 1 and 2 play symmetric roles, thus for example 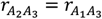, etc.

In Figure 2, we plot the equilibrium value of the important parameter *ρ* for a phenotype with 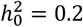 and for different values of the two parameters *r*_*ho*_ and *v* that describe the respective strength of AM and VCT in the population.

**Figure 2:**
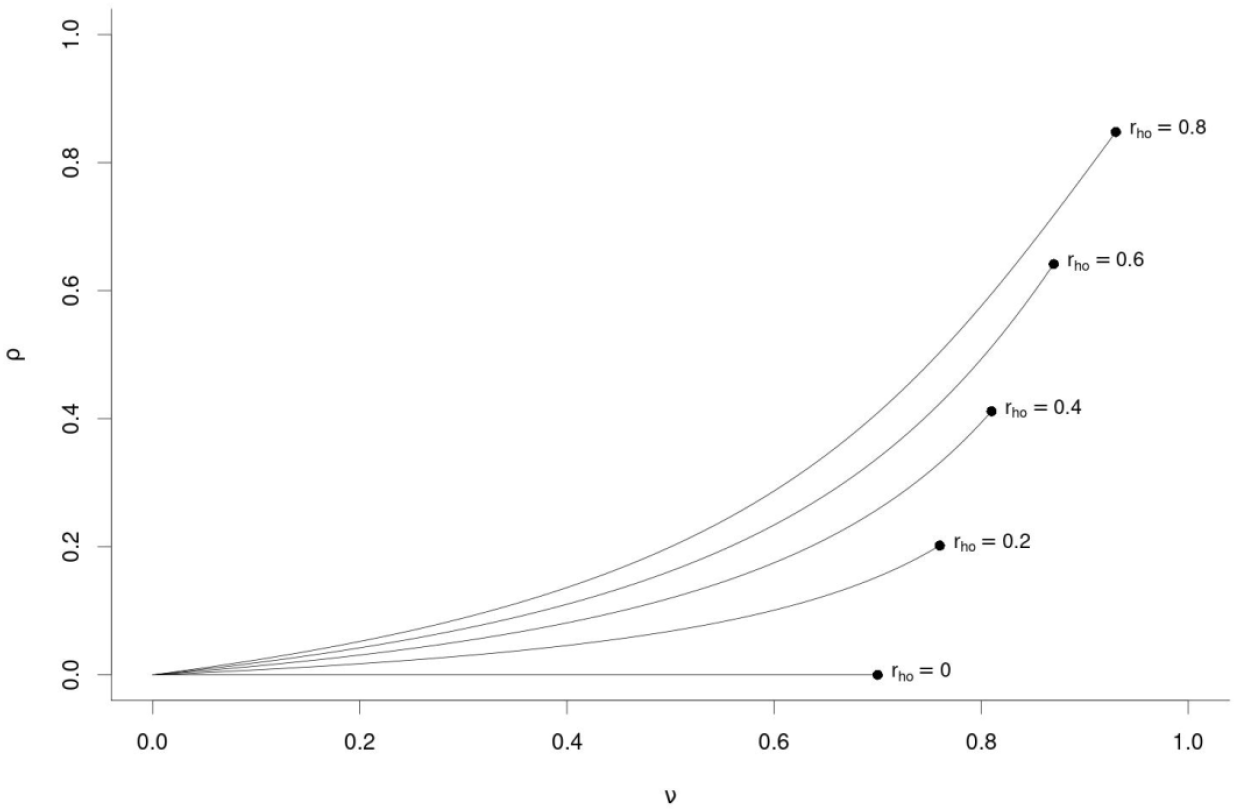
Equilibrium value of *ρ* as a function of *r*_*ho*_ and *v*. Note that there is an upper bound to *v*, depending on *r*_*ho*_ and 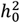. For example, when *r* =0 and for any 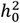, and thus the environments of the mates are independent, *v* cannot be higher that 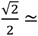 0.7. Higher values of *r*_*ho*_ imply a positive correlation between the mates’ environment, allowing *v* to take higher values.

Furthermore, we show the inflation of estimates of 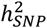 in the presence of co-occurring AM and VCT as a function of *r*_*ho*_ and *v* (Figure 3).

**Figure 3:**
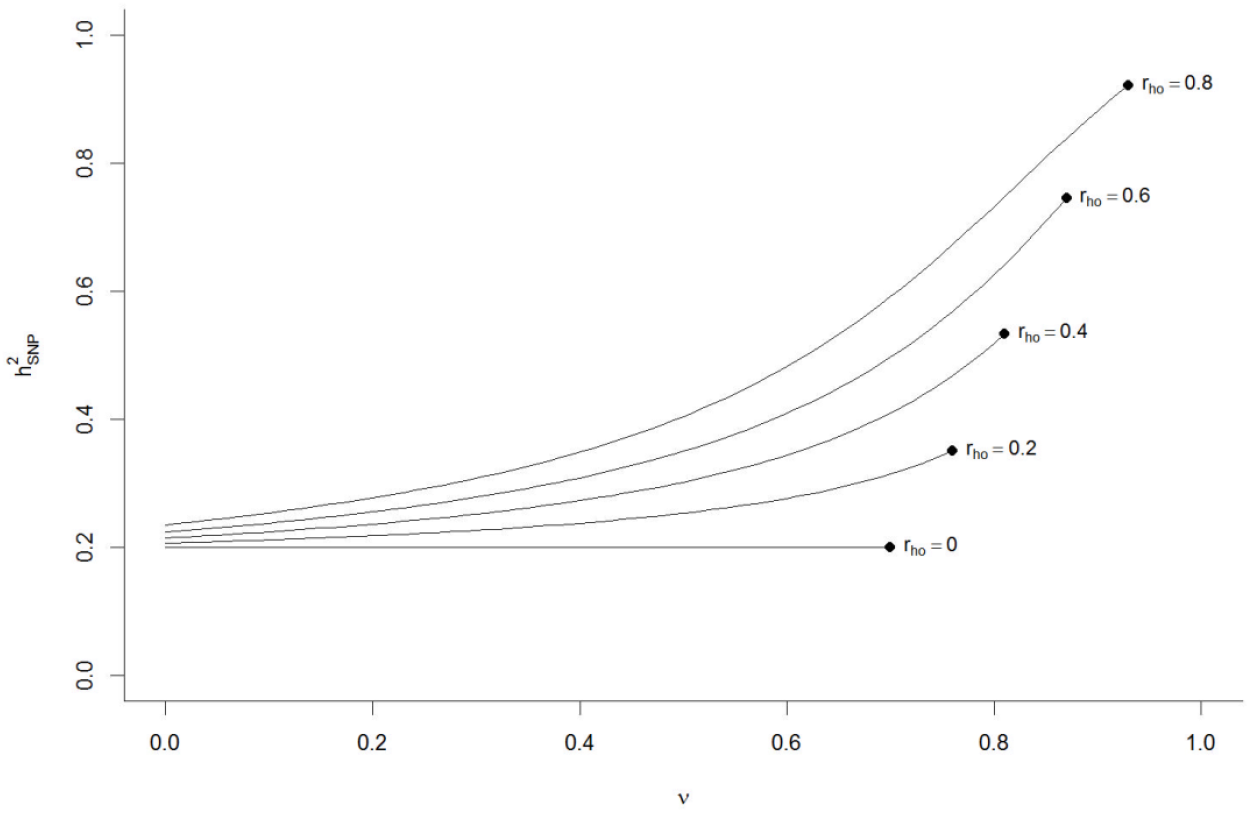
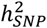 at equilibrium as a function of *r*_*ho*_ and *v* for a phenotype with 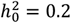 in generation 0 (before AM and VCT).

### 3.2 Consequences on global association statistics (SNP-heritability, GWAS, and Polygenic Scores)

In this section, we examine the impact of the AMVCT model of commonly used analyses in genetic epidemiology. The derivations of these results are given in full in section 6 in the Supplementary Material.

#### 3.2.1 SNP-heritability

Considering pairs of unrelated individuals with a fixed value correlation between their genetic values, say cor(*A* _*i*,1_, A_*i*,2_) =*u*, where the pairs are indexed by *i*, then one computes cor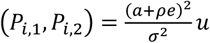.

Then the SNP-heritability, which can be estimated as cor(*P*_*i*,1_, *P*_*i*,2_)/cor(*A*_*i*,1_, *A*_*i*,2_) in the spirit of Haseman-Elston regression, is equal to

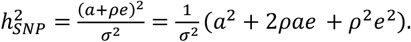

It is worth to note here that to unravel the genetic architecture of the trait, one would need to estimate separately the three phenotype variance components: *a*^2^, 2*ρ ae* and *e*^2^; the value of 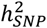 not only includes the two first components, but even a part of the last one (full details in section 6.1 of the Supplementary Materials).

We depict the three components of total phenotypic variance that arise from the presence of the correlation *ρ* induced by the co-occurrence of AM and VCT in Figure 4; showing how estimates of SNP-heritability will be inflated and even capture a small part of the purely environmental variance.

**Figure 4:**
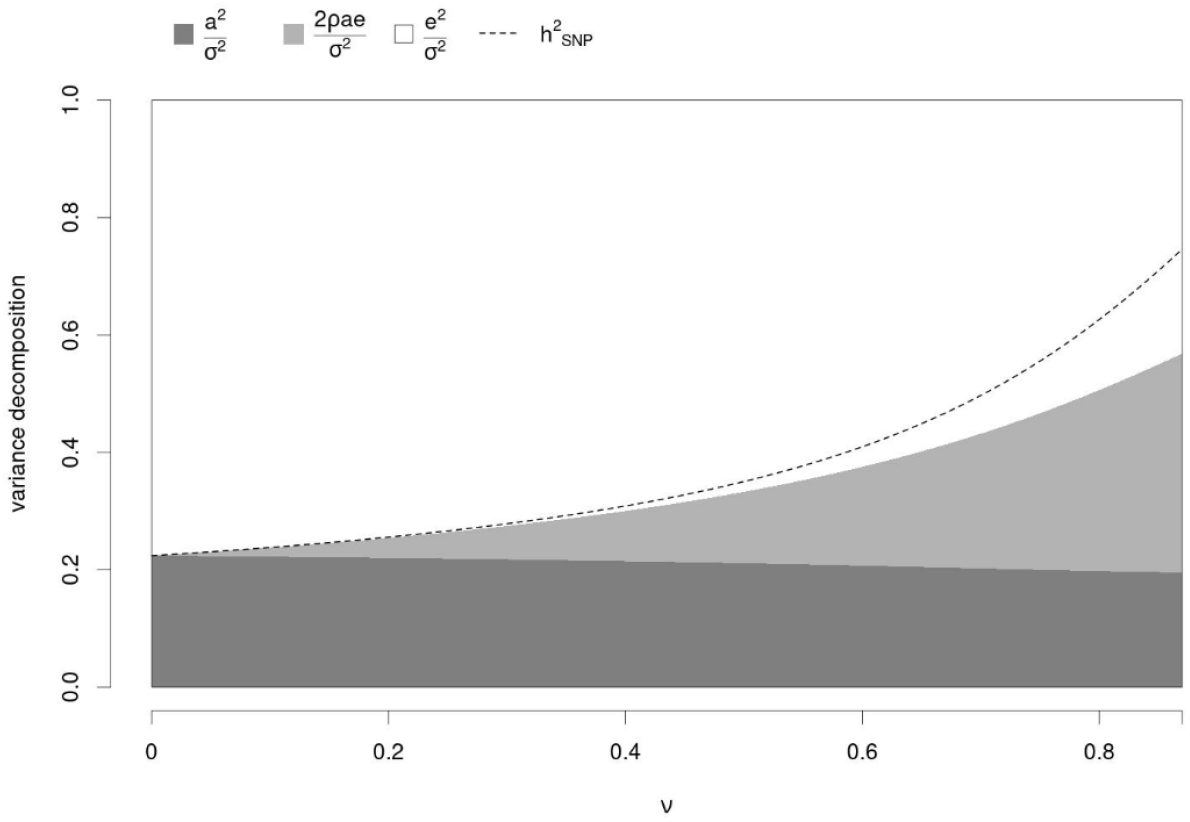
Decomposition of the variance as a function of *v* for a population with heritability 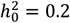 prior to assortative mating; here *r*_*ho*_ =0.6. The dark gray part corresponds to 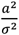, the purely genetic component of the variance; the light gray part to 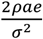, the component created by the positive correlation between genetic and environmental factors; the white part to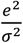, the purely environmental component. The three parts always sum to 1 as seen in the range on the y-axis. The dotted line is the value of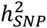, which corresponds to the sum of the two first components, plus a small part of the third.

#### 3.2.2 Consequences on marginal SNP-association statistics

We consider a locus with an allelic substitution effect *β*. The expected value of an estimate 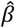 obtained through a linear regression depends on the covariance between the phenotype *P* and the genotype at the locus. This covariance is impacted by the correlation between the causal loci on a given gamete, by the correlation *r*_*ga*_ between the two gametes, and by the correlation *ρ* between the genetic value *A* and the environmental effect *E*. We show in the Supplementary Materials (section 6.2) that 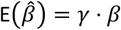 where the inflation factor *γ* is

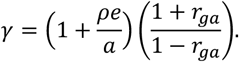

It is worth noting that the multiplicative terms in *γ* all make sense separately: 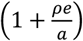 arises due to the gene-environment correlation *ρ*, 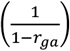 is due to the correlations between causal loci on a same gamete, and 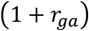 to the correlation between the two gametes. We show the inflation factor gamma in the presence of co-occurring AM and VCT as a function of *r*_*ho*_ and υ (Figure 5).

**Figure 5:**
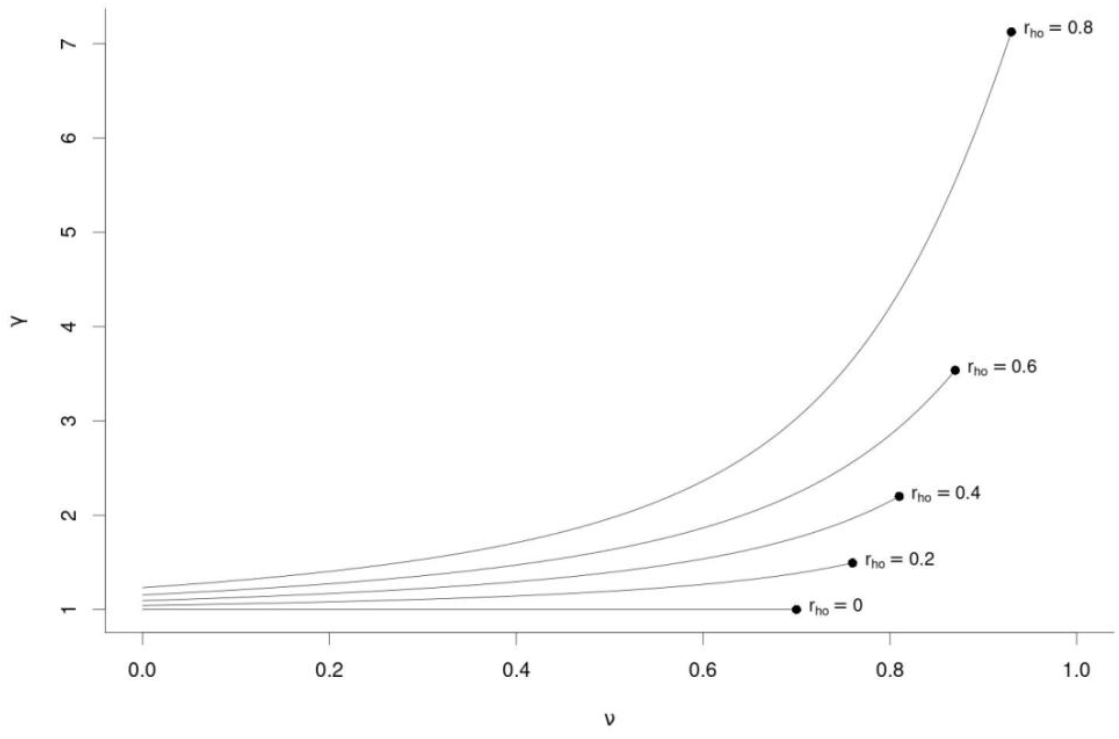
Inflation factor *γ* of SNP effect sizes at equilibrium as a function of *r*_*ho*_ and *v* for a phenotype with 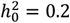 in generation 0 (before AM and VCT).

#### 3.3.3 Polygenic scores

The coefficients 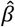 are over-estimated, however the inflation factor *γ* is constant, thus the expected value of the PGS, *Â*, is (leaving aside the question of identifying the causal variants) *E*(*Â*) = *γA*. The proportion of the phenotypic variance explained by the PGS will be (at most) cor(*A, P*)^2^, which is precisely 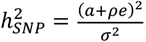.

Of course, one must consider the fact that the values 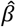 are estimated on a finite sample, so this is an upper bound. As described in (Daetwyler et al., 2008), the PGS would explain less phenotypic variance than 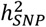 by a factor depending on the size of the discovery sample on which the values 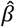 have been estimated as follows:

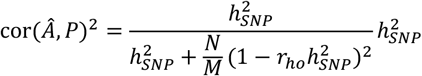

where *M* is the size of the GWAS sample from which the 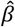 values are estimated. In the context on modern biobanks and large meta-analyses, the ratio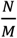 could be argued to be close to 1 or even smaller (Yengo et al., 2022), and as *M* grows very large, this correlation will approach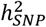. We give details of this calculation, equivalent to (Daetwyler et al., 2008) but under the AMVCT model, in section 6.3 of the Supplementary Materials. Here we do not model the fact that the variants included in the GPS are not in general the causal loci, but loci in tight linkage disequilibrium with those, and that some causal loci may be completely overlooked, which would further shrink the correlation.

#### 3.3.4 Correlation between partial polygenic scores

One can also consider the correlation between PGS computed on different subsets of the genome. Let *S*_1_ and *S*_2_ be two disjoint sets of causal loci (e.g., the loci on even/odd chromosomes). Let *α*_1_ =|*S*_1_|/*N* and *α*_2_ =|*S*_2_|/*N* denote the proportion of causal variants included in these subsets. We show in section 6.4 in Supplementary Materials that the correlation of genetic values 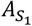and 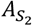 computed on these sets is as follows:

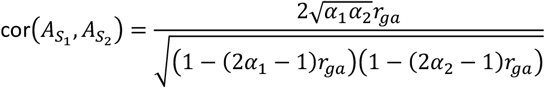

An important case is 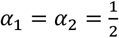, which should hold approximately when one of the sets is make of the odd numbered chromosomes, and the other one of the even numbered chromosomes (Yengo et al., 2018); in this case, one has simply cor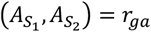. In this particular case, if the partial genetic values are replaced by their estimates from polygenic scores, 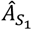 and 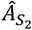, then the correlation becomes:

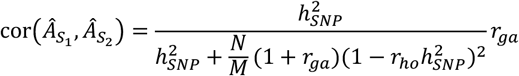

### 3.3 Impact on large-scale family-based association studies

#### 3.3.1 Mid-parent regression

In section 7 of the Supplementary Materials, we investigate further approaches for evaluating genetic architecture using family data. First, we considered estimating heritability via regression of offspring phenotypes on mid-parent phenotypes in trios (Nagylaki, 1978) (section 7.2). This approach gives an unbiased estimate of heritability with a model including only AM; but we show that in the presence of both AM and VCT this gives an inflated estimate of heritability:

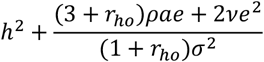

Note that if *v* =0 then *ρ* =0 (see Figure 2) and there is no inflation in the estimate.

#### 3.3.2 Family-based GWAS

In (Howe et al., 2022), the method of sib-GWAS for side-stepping the impact of assortative mating and other GWAS confounders was proposed. The concept is to adjust the regression of phenotype against genotype for the mean parental genotype; essentially looking to capture the true substitution effect of a SNP by examining the difference between genotypes and their expected value (mean parental genotype). As parental genotypes are often unavailable, this method uses a proxy of mean genotypes across sibships, playing on the abundance of sibships in the UK Biobank. A related and more general approach for capturing the ‘direct’ substitution effect of a SNP using parental genotypes can also be found in (A. I. Young et al., 2022). Having shown the inflation of estimates of a GWAS 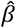 under our framework, we show in section 7 of the Supplementary Materials that adjusting for mean parental genotype (section 7.3) or mean sibling genotype (section 7.4) will indeed produce unbiased estimators of *β*. Hence the shrinkage in effect sizes reported by (Howe et al., 2022) would correspond in our model to the inflation factor *γ*.

We also considered the technique of regressing sibling differences in phenotypes against sibling difference in (estimated) genetic values in section 7.5 of the Supplementary Material; a technique that can be applied (including in the presence of AM) to isolate the role of direct genetic effects from ‘indirect’ genetic effects coming from parental genotypes (Brumpton et al., 2020; A. I. Young et al., 2018). Under our model of AMVCT, we provide the calculation of the variance of sibling phenotype difference explained by difference in their genetic values; as has been previously investigated (Fletcher et al., 2024). Under our model of AMVCT, we arrive at the same result as (Fletcher et al., 2024): indeed, we observe that this technique isolates the direct genetic effects though it does not directly give an estimate of the ‘true heritability’ of the trait due to shared environmental effects. In our model, such shared environmental effects arise solely from VCT and so we parametrise them accordingly in our calculation. We also include the underestimation of the variance component for direct genetic effects in this framework when a previously estimated polygenic score is used rather than the true genetic values of the two siblings.

Finally, in section 7.6 of the Supplementary Material, we consider a further potential study design in family-based GWAS where parental alleles are used to query the ‘Nature of Nurture’ hypothesis (Kong et al., 2018). This proposes that the untransmitted alleles can have an (indirect) effect on the phenotype of the offspring. Here we do not explore the additional complexity that would arise from a model where the offspring phenotype truly has such an ‘indirect genetic’ component. We simply evaluated the expected results of comparing untransmitted alleles to the offspring phenotype under our AMVCT model for a population at equilibrium. In section 7.6 of the Supplementary Material, we first show that the correlation between the ‘untransmitted polygenic score’ in a trio with the offspring phenotype is as follows:

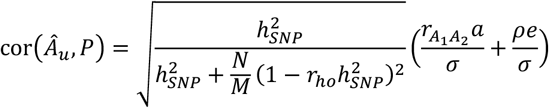

In this formulation, we can interpret the separate contributions from AM in the form of 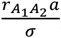, of VCT in the form of 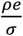, and from the bias in the estimation of the genetic value with a polygenic score coming from the leading radical term; which was already detailed in a previous section above on ‘Polygenic Scores’.

In section 7.6 of the Supplementary Material, we also show that if the untransmitted genotype is regressed against offspring phenotype for causal variant *i* with true allelic substitution effect *β*_*i*_, then the expected regression coefficient of this ‘untransmitted GWAS’ will be 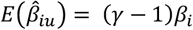. Here, *γ* is the same shrinkage factor as detailed in section 3.2.2. This means that with a large number of trios, one could perform both a standard GWAS in the offspring and an ‘untransmitted allele’ GWAS and under our AMVCT model, the difference in the regression coefficients would have expected value 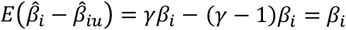; thus this provides another route to an unbiased estimate of the true allelic substitution effect under our model.

### 3.4 Forward-time Simulations

Using our simulated data (see section 2.2), we measured the observed correlation between genetic values and environmental values (i.e. *A*_3_ and *E*_3_) and compared it to the theoretical expectation for *ρ* until the 15^*th*^ generation for populations of size *M* =25,000 and *M* =100,000 (Figure 6). Results show good concordance between the observed and the expected correlation across generations, for both population sizes.

**Figure 6:**
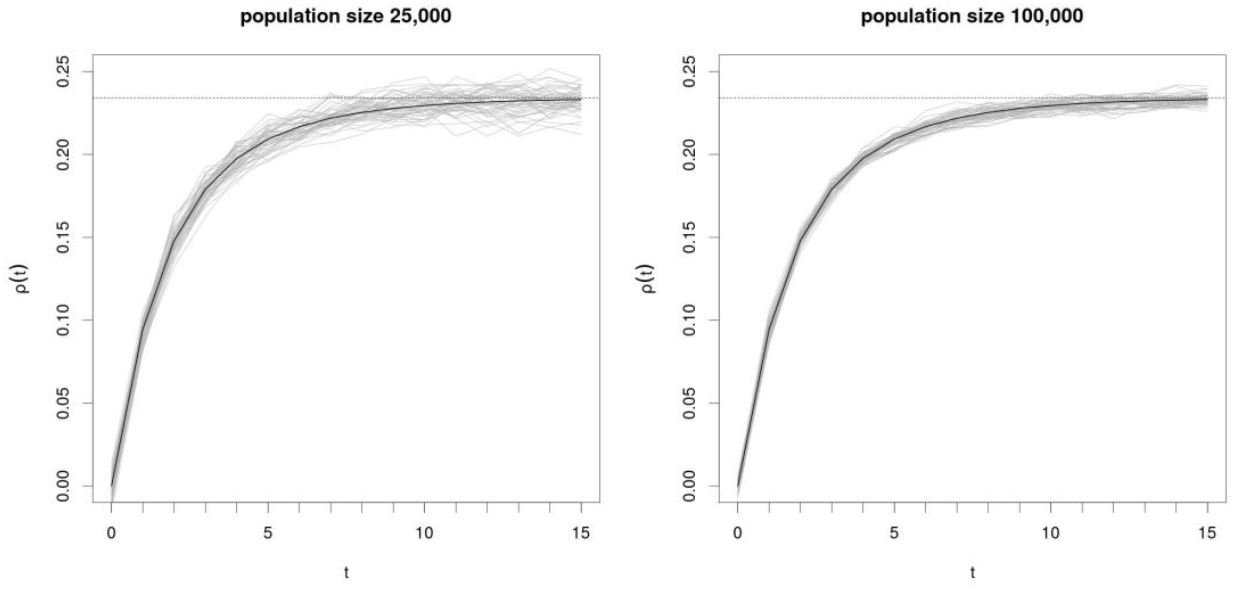
Simulation of the evolution of *ρ*, generations: *t* against observed *ρ* in generation *t*: *ρ*(*t*). Dark line corresponds to the theoretical derivation of the evolution of *ρ*, the dotted line corresponds to the theoretical equilibrium point. Each light grey line shows the observed evolution of *ρ* in a single simulation run (50 simulation runs plotted). AM was simulated with *r*_*ho*_ =0.6 and VCT with *v* =0.6.

Our predictions also held for other parameters of interest (Figure 7).

**Figure 7:**
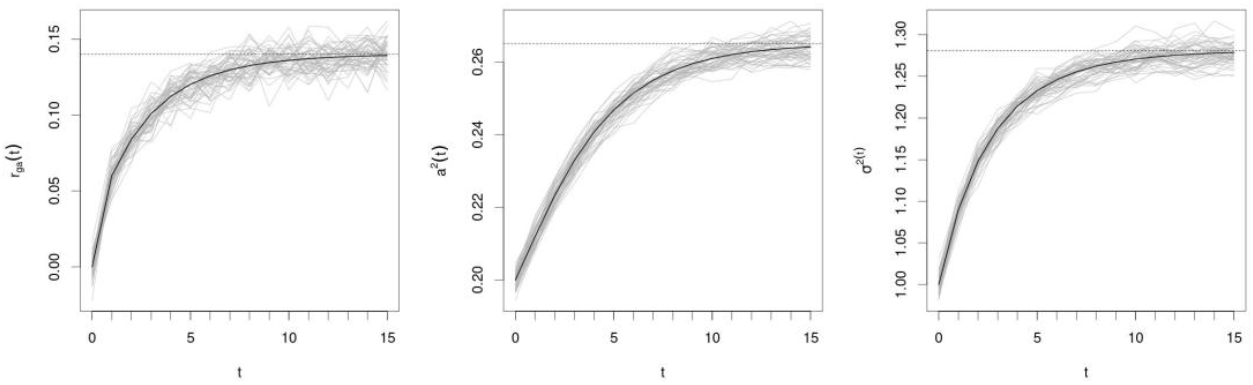
Evolution plots of a similar fashion to Figure 6 for the parameters *r*_*ga*_, *a*^2^, and *σ*^2^. Here, only a population of size *M* =25,000 is shown.

## 4 Discussion

Here we have explored the co-occurrence of two aspects of population architecture that have previously been shown to greatly impact genetic epidemiological studies in human populations: assortative mating and vertical cultural transmission under a classical polygenic additive model. We have set down the formal interplay of these two phenomena, crucially showing an emergent correlation *ρ* between genetic and non-genetic factors that leads to inflation of both heritability and genotype-phenotype association statistics.

After rising sharply for a small number of generations of both AM and VCT, all parameters eventually converge to the theoretical equilibrium points in roughly 15 generations. Through simulations, it was possible to confirm both our calculations of the equations of the evolution and the equilibrium values of *ρ* as well as other quantities of interest.

We were also able to evaluate the impact on estimations coming from population-based studies including inflations to estimates of SNP-heritability, GWAS effect sizes and hence polygenic scores. Whilst it may not be possible to attain a precise model for mate-choice or for the correlation of environmental factors, the huge breadth of recent genetic cross-sectional studies should allow for more nuanced approaches than assuming panmixia and completely random environmental effects.

The search for traces for assortative mating in genetic datasets has largely concentrated on observing long range correlation between variants detected through GWAS, notably using techniques such as observing genetic correlation between odd and even chromosomes (Yengo et al., 2018). An alternative method for observing genetic traces of AM in population-based studies could be to directly calculate gametic correlation though it is normally not possible to determine the required parent-of-origin information. However, recent advances in parent-of-origin determination in biobank-type datasets (Hofmeister et al., 2022) using ideas of surrogate parent groups (Kong et al., 2008) could allow such an approach to become practical.

The crucial observation in this work is that even in the absence of all confounders such as population/cultural stratification or ascertainment bias, genetic associations can still capture environmental factors. This would clearly add to the numerous difficulties in interpreting many GWAS associations and the predictive performance of PGS (Janssens, 2019; Pingault et al., 2022). In particular, if an estimated PGS is capturing a part of the environment effects, using the PGS as a covariable in order to investigate environmental effects, as discussed in (Uddin et al., 2022), would not be appropriate. Furthermore, whilst the impact of VCT and AM on Mendelian Randomization (MR) have previously been studied with methods for bias adjustment (Brumpton et al., 2020; Davies et al., 2019; Hartwig et al., 2018; Howe et al., 2019), their co-occurrence could be particularly disruptive for MR studies.

We confirmed that analysing family-data from trios or quads can provide unbiased estimates of allelic substitution effects in the presence of co-occurring AM and VCT; though we have not explored more complicated models for shared environmental effects in families beyond a simple model of positive covariance between environments between generations. In this work, we did not model ‘indirect’ genetic effects, where parental phenotypes (via parental genotypes) affect the phenotypes of the offspring; an in-depth exploration of this model in the presence of AM is detailed in (A. S. Young, 2023). We have however shown how the AMVCT model will lead to correlations between untransmitted alleles (in the form of single variants or as polygenic scores) with offspring phenotypes which could be interpreted as evidence of indirect genetic effects of parental genotypes; whereas these correlations arise in our model solely from the correlations between parental genetic values and between genetic and environmental components within an individual.

## 5 Conclusions

The list of strong model assumptions that underlie the classic additive polygenic model is extensive, of which random mating and random environmental factors are just two. We have set out a framework for the co-occurrence of AM and VCT, but it must be noted that non-random mating would be unlikely to depend on a single trait and be independent of population/cultural stratification. Such stratification could also reflect differences in the degree of VCT and so we have likely only scratched the surface of one of the processes that leads to genetic and non-genetic factors become correlated in human populations.

## Author Contributions

HP and AFH derived the theoretical framework and performed the analyses. All authors participated in the review of the literature, the design of our study and the writing of the manuscript.

## Data availability

Scripts for replicating the simulation studies presented in this work and for computing the evolution and equilibrium values of all parameters under the AMVCT model are available at https://github.com/HervePerdry/AMVCTpaper. No real human genetic data was used in this study.

