## Supplementary Materials for "Impact of co-occurrent assortative mating and vertical cultural transmission on measures of genetic associations"

[1]Univ. Brest, Inserm, EFS, UMR 1078, GGB, Brest, France
[2]Laboratoire Cogitamus
[3]CESP Inserm, U1018, Université Paris-Saclay, UVSQ, Villejuif, France.

**References** **41**

**A Technical lemma** **42**

**B Computing the evolution of a population** **45**

### Introduction

This is the supplementary material to ”Impact of co-occurrent assortative mating and vertical cultural transmission on measures of genetic associations”.

In the first section below, we introduce the polygenic additive model, and the notations used in the whole document.

In the second section, we consider the case of a population with assortative mating (AM), and derive correlations between genetic and environmental components of any two mates.

In the third section, we consider offsprings in such a population, and we introduce our model for vertical cultural transmission (VCT). We derive the equation of evolution of gametic correlation and gene-environment correlation between generation $t$ and $t+1$.

In the fourth section, we consider the case were all causal loci effects have the same variance. In this case, we derive the evolution of the inter-loci correlations between times $t$ and $t+1$, and show how to compute numerically the value of all quantities of interest of the model.

In the fifth section, we consider the general case where causal loci effects have different variances, and show that the conclusions of the fourth section still hold.

In the sixth section, we investigate the behaviour of classical analyses employed in genetic epidemiology (estimation of SNP-heritability, association analysis, polygenic scores) in the presence of assortative mating and vertical cultural transmission.

In the seventh section, we extend by examining the impact of assortative mating and vertical cultural transmission on family-based study designs.

### 1 The additive polygenic model

#### 1.1 Notations

We consider a phenotype determined by $P = A + E$, where $A$ is the sum of independent genetic effects, and $E$ is the environmental effect. We denote $\mathrm{var}(P) = \sigma^2$, $\mathrm{var}(A) = a^2$ and $\mathrm{var}(E) = e^2$. In presence of a gene-environment correlation $\rho = \mathrm{cor}(A, E)$, one has $\sigma^2 = a^2 + 2\rho ae + e^2$.

In parts of what follows, $a^2, \rho$ and $\sigma^2$ will depend on the generation $t$ ; they will later be denoted $a^2(t)$, $\rho(t)$ and $\sigma^2(t)$. In this section and the following, we are at fixed time $t$. The environmental variance $e^2$ will be considered as constant through time.

In the sequel, we will introduce notations for several quantities that are equal to zero in a random mating population, but will prove to be non-zero in the random mating with cultural transmission model, e.g. the correlation between the parental gametes.

#### 1.2 Loci and gametes

We consider $N$ causal di-allelic loci, indexed by $i = 1, \dots, N$. Throughout what follows, $N$ is assumed to be very large.

In a given indidual's genome, the alleles at these loci are carried by a paternal gamete $G^p$ and a maternal gamete $G^m$. We let $X_i^p$ and $X_i^m$ be identically distributed centered random variables encoding these alleles: if $p_i$ is the frequency of the reference allele and $q_i = 1 - p_i$ is the frequency of the alternate allele, we have $X_i^p = -q_i$ (for the reference allele) or $X_i^p = p_i$ (for the alternate allele), with respective probability $p_i$ and $q_i$. We have $E(X_i^p) = 0$ and $\text{var}(X_i^p) = p_i q_i$. The genotype of the individual is encoded by $X_i = X_i^p + X_i^m$. We have $E(X_i) = 0$.

Let $\beta_i$ be the allelic effect of locus $i$ on the phenotype $P$. We let $g_i^p = \beta_i X_i^p$ and $g_i^m = \beta_i X_i^m$. These two variables are identically distributed, with expected value 0 and variance $\beta_i^2 p_i q_i$. The genetic value of the paternal (respectively maternal) gamete is $G^p = \sum_{i=1}^N g_i^p$ (respectively $G^m = \sum_{i=1}^N g_i^m$).

The (additive) genetic value of the individual is $A = G^p + G^m$. Letting $g_i = g_i^p + g_i^m = \beta_i X_i$, we have also $A = \sum_{i=1}^N g_i$.

As $N$ is large, the distribution of $G^p$, $G^m$, and $A$ can be considered as Gaussian.

##### 1.3 Inter-loci correlation

If the variables $g_i^p$ are independent, then the variance of the genetic value of a gamete is

$$g_0^2 = \text{var}(G^p) = \sum_{i=1}^N \beta_i^2 p_i q_i \tag{1}$$

We are going to introduce notations allowing to express $\text{var}(G) = g^2$ in the general case. Let

$$\kappa_i^2 = N \frac{\beta_i^2 p_i q_i}{g_0^2}, \tag{2}$$

so that

$$\text{var}\big(g_i^p\big) = \frac{1}{N} g_0^2 \kappa_i^2. \tag{3}$$

From equations 1 and 2, we have $\sum_{i=1}^N \kappa_i^2 = N$. If all loci contribute to the genetic value with the same variance, we will have $\kappa_i^2 = 1$ for all $i$. In general, the $\kappa_i$ are assumed to be close to one (even if the loci contribute with different variances, they have same order of magnitude).

We don't assume that the loci carried by a gamete are independent. Let $\kappa_{ij}$ be such that

$$\text{cov}\big(g_i^p, g_j^p\big) = \frac{1}{N} g_0^2 \kappa_{ij}. \tag{4}$$

Thus, $\kappa_{ii} = \kappa_i^2$ and

$$\text{cor}\big(g_i^p, g_j^p\big) = \frac{\kappa_{ij}}{\kappa_i \kappa_j}. \tag{5}$$

Then

$$\begin{aligned} \text{var}(G^p) &= \sum_{i,j} \text{cov}\big(g_i^p, g_j^p\big) \\ g^2 &= \frac{1}{N} g_0^2 \sum_{i,j} \kappa_{ij} \\ g^2 &= N g_0^2 \overline{\kappa} \end{aligned} \tag{6}$$

where

$$\overline{\kappa} = \frac{1}{N^2} \sum_{i,j} \kappa_{ij}.$$

Note that $g^2$ and $g_0^2$ should have the same order of magnitude, thus $\overline{\kappa}$ has to be infinitesimal (it has order of magnitude $\frac{1}{N}$), and the $\kappa_{ij}$ as well.

#### 1.4 Gametic correlation

We denote $G^p$ and $G^m$ the genetic value of the parental gametes carried by an individual. We assume between these values a gametic correlation $r_{\text{ga}} = \text{cor}(G^p, G^m)$. The variance of $A = G^p + G^m$ is

$$\text{var}(A) = a^2 = 2(1 + r_{\text{ga}})g^2. \tag{7}$$

#### 1.5 Correlation between loci and gamete values

As $G^p = \sum_i g_i^p$, we have

$$\text{cov}\big(g_i^p, G^p\big) = \frac{1}{N} g_0^2 \sum_i \kappa_{ij} = g_0^2 \overline{\kappa}_i, \tag{8}$$

where $\overline{\kappa}_i = \frac{1}{N} \sum_i \kappa_{ij}$.

The correlation is then

$$\text{cor}\big(g_i^p, G^p\big) = \frac{g_0^2 \overline{\kappa}_i}{\sqrt{\frac{1}{N} g_0^2 \kappa_i^2} \cdot \sqrt{N g_0^2 \overline{\kappa}}},$$

and finally

$$\text{cor}\big(g_i^p, G^p\big) = \frac{\overline{\kappa}_i}{\kappa_i \sqrt{\overline{\kappa}}}. \tag{9}$$

#### 1.6 Meiosis

Consider an individual with genetic component $A = G^p + G^m$, with $G^p = \sum_{i=1}^N g_i^p$ and $G^m = \sum_{i=1}^N g_i^m$. To model the genetic value of a gamete emitted through meiosis by this individual, we let $I_p \cup I_m = [1, \dots, N]$ with $I_p \cap I_m = \varnothing$ be the sets of indices of the transmitted alleles of paternal and maternal origins respectively. Each $i$ is in $I_p$ with probability $\frac{1}{2}$ so in practice $I_p$ and $I_m$ have similar size and play symmetric roles.

The transmitted gamete has genotypes $X_i^p$ if $i \in I_p$, and $X_i^m$ if $i \in I_m$. Its genetic value is

$$G = \sum_{i \in I_p} g_i^p + \sum_{i \in I_m} g_i^m.$$

By letting

$$H = \sum_{i \in I_p} g_i^m + \sum_{i \in I_m} g_i^p,$$

one gets $A = G + H$; $H$ can be seen as the genetic value of the “untransmitted gamete”.

The gametic variance is $\mathrm{var}(G) = \mathrm{var}(H) = g^2$. We have $a^2 = \mathrm{var}(G+H) = 2g^2 + \mathrm{cov}(G,H)$, thus $\mathrm{cov}(G,H) = \frac{1}{2}a^2 - g^2 = g^2(1+r_{\mathrm{ga}}) - g^2 = g^2 r_{\mathrm{ga}}$, hence the correlation between $G$ and $H$ is

$$\mathrm{cor}(G,H) = r_{\mathrm{ga}}. \tag{10}$$

Moreover, one has $\mathrm{cov}(A,G) = \mathrm{cov}(G+H,G) = g^2 + g^2 r_{\mathrm{ga}}$, thus (cf equation 7)

$$\mathrm{cov}(A,G) = (1+r_{\mathrm{ga}})g^2 = \frac{1}{2}a^2. \tag{11}$$

Using the properties of multivariate Gaussian distributions, one gets that the distribution of $G$ conditional to $A$ is Gaussian, with expected value $\frac{1}{2}A$ and variance $\frac{1}{2}(1-r_{\mathrm{ga}})g^2$:

$$G \mid A \sim \mathcal{N}\left(\frac{1}{2}A, \frac{1}{2}(1-r_{\mathrm{ga}})g^2\right) \tag{12}$$

### 2 Individuals and mate pairs in a population with assortative mating

The results of this section are for a fixed time $t$. To simplify notations, we don't write $g(t)$, $a(t)$, $\sigma^2(t)$, but $g$, $a$, $\sigma^2$, etc. The phenotype of one individual is $P = A + E$, with the genetic value $A$ having a variance $\mathrm{var}(A) = a^2$ and the variance of environmental contribution is $\mathrm{var}(E) = e^2$. We assume there can be some gene-environment correlation $\rho = \mathrm{cor}(A,E) = r_{AE}$.

Consider two mates, indexed by 1 and 2, with phenotypes $P_1 = A_1 + E_1$ and $P_2 = A_2 + E_2$. We assume that there is some assortative mating in the population, modeled by a single parameter $r_{\mathrm{ho}} = \mathrm{cor}(P_1, P_2) = r_{P_1P_2}$.

We will prove that all four variables $A_1$, $E_1$, $A_2$ and $E_2$ are correlated, with correlations given by the following table (for the sake of readibility, we don't include the bottom part of the correlation matrix which is symmetric, nor its diagonal which is equal to 1).

| | $E_1$ | $A_2$ | $E_2$ |
|---|---|---|---|
| $A_1$ | $\rho$ | $\frac{r_{\mathrm{ho}}}{\sigma^2}(a+\rho e)^2$ | $\frac{r_{\mathrm{ho}}}{\sigma^2}(\rho a + e)(\rho e + a)$ |
| $E_1$ | | $\frac{r_{\mathrm{ho}}}{\sigma^2}(\rho a + e)(\rho e + a)$ | $\frac{r_{\mathrm{ho}}}{\sigma^2}(\rho a + e)^2$ |
| $A_2$ | | | $\rho$ |

**Table 1:** Correlations between $A$ and $E$ terms for the two parents

#### 2.1 A single individual in the population

We first derive expressions for the correlations $r_{AP}$ and $r_{EP}$ that will be useful later.

We have

$$\mathrm{var}(P) = \sigma^2 = a^2 + 2\rho a e + e^2,$$

and

$$\begin{aligned}\mathrm{cov}(A,P) &= \mathrm{cov}(A, A+E) \\ &= \mathrm{var}(A) + \mathrm{cov}(A,E) \\ &= a^2 + \rho a e\end{aligned}$$

that is

$$\operatorname{cov}(A,P) = a(a+\rho e) \tag{13}$$

and

$$r_{AP} = \frac{1}{\sigma}(a+\rho e) \tag{14}$$

In the same way,

$$\begin{aligned}\operatorname{cov}(E,P) &= \operatorname{cov}(E, A+E)\\ &= \operatorname{cov}(A,E) + \operatorname{var}(E)\\ &= \rho a e + e^2\end{aligned}$$

and

$$r_{EP} = \frac{1}{\sigma}(\rho a+e) \tag{15}$$

#### 2.2 A couple with assortative mating

In this section and the next one, we use the same computational approach than Nagylaki (1978) [4].

Consider a couple with two mates indexed by 1 and 2. The values of $P$, $A$ and $E$ for these individuals are denoted $P_i$, $A_i$, $E_i$ ($i = 1,2$).

We assume that mate choice depends on the value of $P$, so that there is a correlation $r_{\text{ho}} = r_{P_1P_2}$ between the two members of a couple. It is assumed that the mate choice is based only on the phenotype, i.e. that conditional to $P_2$, $P_1$ and $A_2$ are independent: $r_{P_1A_2\cdot P_2} = 0$, where $r_{P_1A_2\cdot P_2}$ denotes the correlation between $P_1$ and $A_2$ conditional to $P_2$, and also $r_{A_1A_2\cdot P_2} = 0$, $r_{P_1E_2\cdot P_2} = 0$, $r_{E_1E_2\cdot P_2} = 0$.

##### 2.2.1 Computing $r_{A_1A_2}$

Using lemma 1 (appendix A) with $r_{P_1A_2\cdot P_2} = 0$, we have

$$\operatorname{cor}(P_1, A_2) = r_{P_1A_2} = r_{P_1P_2} r_{A_2P_2}.$$

We have $r_{P_1P_2} = r_{\text{ho}}$ and (equation 14) $r_{A_2P_2} = \frac{1}{\sigma}(a+\rho e)$, hence

$$r_{P_1A_2} = \frac{r_{\text{ho}}}{\sigma}(a+\rho e).$$

The two members of the couple play symmetric roles so one has also $r_{A_1P_2} = \frac{r_{\text{ho}}}{\sigma}(a+\rho e)$.

Applying again lemma 1 with $r_{A_1A_2\cdot P_2} = 0$, we have (using equation 14)

$$r_{A_1A_2} = r_{A_1P_2} r_{A_2P_2} = \frac{r_{\text{ho}}}{\sigma^2}(a+\rho e)^2. \tag{16}$$

##### 2.2.2 Computing $r_{E_1E_2}$

Using again lemma 1 with $r_{P_1E_2\cdot P_2} = 0$, and using equation 15, we get

$$r_{P_1E_2} = r_{P_1P_2} r_{E_2P_2} = \frac{r_{\text{ho}}}{\sigma}(\rho a + e). \tag{17}$$

And by symmetry, we have $r_{E_1P_2} = \frac{r_{\text{ho}}}{\sigma}(\rho a + e)$.

Finally from $r_{E_1E_2\cdot P_2} = 0$, we get (using equation 15)

$$r_{E_1E_2} = r_{E_1P_2} r_{E_2P_2} = \frac{r_{\text{ho}}}{\sigma^2}(\rho a + e)^2. \tag{18}$$

##### 2.2.3 Computing $r_{A_1E_2}$ and $r_{E_1A_2}$

We have, using equations 17 and 18,

$$\begin{aligned}
\operatorname{cov}(P_1, E_2) &= \operatorname{cov}(A_1 + E_1, E_2) \\
r_{P_1E_2} e\sigma &= \operatorname{cov}(A_1, E_2) + \operatorname{cov}(E_1, E_2) \\
r_{\text{ho}}(\rho a + e)e &= r_{A_1E_2} ae + r_{E_1E_2} e^2 \\
r_{\text{ho}}(\rho a + e) &= r_{A_1E_2} a + \frac{r_{\text{ho}}}{\sigma^2}(\rho a + e)^2 e
\end{aligned}$$

hence

$$\begin{aligned}
r_{A_1E_2} a &= \frac{1}{\sigma^2} r_{\text{ho}}(\rho a + e)(\sigma^2 - (\rho a + e)e) \\
&= \frac{1}{\sigma^2} r_{\text{ho}}(\rho a + e)(\rho a e + a^2).
\end{aligned}$$

By symmetry of the roles of the two parents, we have $r_{E_1A_2} = r_{A_1E_2}$, that is

$$r_{E_1A_2} = r_{A_1E_2} = \frac{r_{\text{ho}}}{\sigma^2}(\rho a + e)(\rho e + a) \tag{19}$$

We have proved all the correlations of table 1.

### 3 Offsprings in a population with assortative mating

Here we consider a couple at generation $t$, and their offspring at generation $t+1$. The mates are indexed by $i = 1, 2$ as above, and the offspring has index 3.

We assume that there is a correlation

$$v = r_{E_1E_3} = r_{E_2E_3} \tag{20}$$

between the parental and offspring environment, which can be attributed to cultural transmission.

We will derive some equations of evolution of the parameters of the model between time $t$ and $t+1$, and prove that at equilibrium the correlations between the parents and the offspring are as given in the following table.

| | $A_3$ | $E_3$ |
|---|---|---|
| $A_1$ or $A_2$ | $\frac{1}{2}\left(1+\frac{r_{\text{ho}}}{\sigma^2}(a+\rho e)^2\right)$ | $\rho$ |
| $E_1$ or $E_2$ | $\frac{1}{2}\left(\rho+\frac{r_{\text{ho}}}{\sigma^2}(a+\rho e)(\rho a+e)\right)$ | $\nu$ |

**Table 2:** Correlations at equilibrium between $A$ and $E$ terms for parent / offspring pairs.

We will first establish equations of evolutions for $r_{\text{ga}}(t)$, $a(t)$, $r_{A_1A_3}(t)$, $r_{A_1E_3}(t)$ and $\rho(t)$. The equilibrium values of $r_{A_1A_3}$, $r_{A_1A_3}$, and $r_{A_1E_3}$ in the above table will then be deduced from these equations. However the computation of the equilibrium value of $\rho$ and $a$ will be postponed to sections 4 and 5.

##### 3.1 Notations

We assume that the individuals $i=1,2$ emit gametes with genetic values $G_i$, with the genetic value of the individual decomposing in $A_i = G_i + H_i$, as in subsection 1.6, where $G_i$ is the genetic value gamete transmitted to the offspring and $H_i$ is the genetic value of the "untransmitted gamete". The genetic value of the offspring is $A_3 = G_1 + G_2$ (one can also write $A_3 = G_3^p + G_3^m$ with $G_3^p = G_1$, $G_3^m = G_2$).

The individuals $A_1$ and $A_2$ pertain to the generation $t$, and $A_3$ pertains to the generation $t+1$, thus

$$\text{var}(A_1) = \text{var}(A_2) = a^2(t) = 2(1+r_{\text{ga}}(t))g^2(t),$$

and

$$\text{var}(A_3) = a^2(t+1) = 2(1+r_{\text{ga}}(t+1))g^2(t+1). \tag{21}$$

Simarly, we have $\text{cor}(A_3, E_3) = \rho(t+1)$, and

$$\text{cov}(A_3, E_3) = \rho(t+1)a(t+1)e. \tag{22}$$

In the sequel, we will use several times the fact the parents play symmetric roles, thus, for example, $r_{A_2A_3}(t) = r_{A_1A_3}(t)$.

##### 3.2 Equation of evolution of the gametic correlation $r_{\text{ga}}(t)$

We will link the gametic correlation $r_{\text{ga}}(t+1) = \text{cor}(G_1, G_2)$ with $\text{cor}(A_1, A_2) = r_{A_1A_2}(t)$.

We write

$$\text{cov}(A_1, A_2) = \text{cov}(G_1, G_2) + \text{cov}(G_1, H_2) + \text{cov}(H_1, G_2) + \text{cov}(H_1, H_2).$$

All four terms on the right hand side are equal, and $\text{cov}(A_1, A_2) = r_{A_1A_2}(t)a^2(t)$, so

$$\text{cov}(G_1, G_2) = \frac{1}{4} r_{A_1A_2}(t)a^2(t).$$

One has $a^2(t) = 2(1+r_{\text{ga}}(t))g^2(t)$ (equation 7) and $\text{cov}(G_1, G_2) = r_{\text{ga}}(t+1)g^2(t+1)$, hence

$$r_{\text{ga}}(t+1) = \frac{1}{2} r_{A_1A_2}(t) \frac{g^2(t)}{g^2(t+1)}(1+r_{\text{ga}}(t)). \tag{23}$$

##### 3.3 Equation of evolution of $r_{A_1A_3}(t)$

We compute hereafter the correlations $r_{A_1A_3}(t)$ and $r_{A_2A_3}(t)$ between the parental genetic components and the offspring's one.

We have (equation 11) $\mathrm{cov}(A_1,G_1) = \frac{1}{2}a^2(t)$. From $\mathrm{cov}(A_1,A_2) = \mathrm{cov}(A_1,G_2)+\mathrm{cov}(A_1,H_2)$ and $\mathrm{cov}(A_1,G_2) = \mathrm{cov}(A_1,H_2)$ (as $G_2$ and $H_2$ play symmetric roles), we get

$$\mathrm{cov}(A_1,G_2) = \frac{1}{2}\mathrm{cov}(A_1,A_2) = \frac{1}{2}r_{A_1A_2}(t)a^2(t).$$

Hence

$$\mathrm{cov}(A_1,A_3) = \mathrm{cov}(A_1,G_1)+\mathrm{cov}(A_1,G_2) = \frac{1}{2}(1+r_{A_1A_2}(t))a^2(t)$$

and

$$r_{A_1A_3}(t) = \frac{\frac{1}{2}(1+r_{A_1A_2}(t))a^2(t)}{a(t)a(t+1)}$$

so

$$r_{A_1A_3}(t) = \frac{1}{2}(1+r_{A_1A_2}(t))\frac{a(t)}{a(t+1)}. \tag{24}$$

As noted before, the two parents play symmetric roles and $r_{A_2A_3}(t) = r_{A_1A_3}(t)$.

##### 3.4 The correlations between parent and offspring environmental components

We have denoted by $v$ the correlation between the offspring's environment $E_3$, and the parental environments $E_1$ and $E_2$ (equation 3.5). All these variables have expected value 0, and $\mathrm{var}(E_i) = e^2$ for $i = 1,2,3$.

The variance-covariance matrix of $(E_1,E_2,E_3)$ is

$$e^2\begin{pmatrix} 1 & r_{E_1E_2} & v \\ r_{E_1E_2} & 1 & v \\ v & v & 1 \end{pmatrix} \tag{25}$$

Moreover, conditionally to $(E_1,E_2)$, $E_3$ is independent from $A_1$ and $A_2$.

Note that $v$ is upper bounded; for example if $r_{E_1E_2} = 0$, the maximum value of $v$ in that case is $v = \frac{1}{\sqrt{2}}$, in which case $E_3$ is fully determined by $E_1$ and $E_2$, and we have $E_3 = \frac{1}{2}(E_1+E_2)$. In general, the above matrix is positive definite if

$$v \le \sqrt{\frac{1+r_{E_1E_2}}{2}}. \tag{26}$$

The value $v = \sqrt{\frac{1+r_{E_1E_2}}{2}}$ corresponds to the extreme case in which the offspring's environment is entirely determined by its parents' environments, that is again $E_3 = \frac{1}{2}(E_1+E_2)$.

##### 3.5 Equation of evolution of the correlation $r_{A_1E_3}(t)$

We want the equation of evolution of the correlation between the parental genetic value $A_1$ (or $A_2$) and the offspring environment $E_3$.

Let us compute $\text{cov}(E_1, A_3) = \text{cov}(E_1, G_1) + \text{cov}(E_1, G_2)$. We have $\text{cov}(E_1, A_1) = \text{cov}(E_1, G_1) + \text{cov}(E_1, H_1)$ ; the two terms in the right hand side are equal, so $\text{cov}(E_1, G_1) = \frac{1}{2}\text{cov}(E_1, A_1)$. In the same way, $\text{cov}(E_1, G_2) = \frac{1}{2}\text{cov}(E_1, A_2)$. Hence

$$\begin{aligned}\text{cov}(E_1, A_3) &= \text{cov}(E_1, G_1) + \text{cov}(E_2, G_2) \\ &= \frac{1}{2}\text{cov}(E_1, A_1) + \frac{1}{2}\text{cov}(E_1, A_2) \\ &= \frac{1}{2}\rho(t)ea(t) + r_{E_1 A_2}(t)ea(t) \\ &= \frac{1}{2}\left(\rho(t) + \frac{r_{\text{ho}}}{\sigma^2(t)}\big(\rho(t)a(t)+e\big)\big(a(t)+\rho(t)e\big)\right)ea(t),\end{aligned}$$

where the value of $r_{E_1 A_2}$ is taken from table 1 (and equation 19).

One has $\text{cov}(A_3, E_3) = \text{cov}(G_1, E_3) + \text{cov}(G_2, E_3) = \frac{1}{2}\text{cov}(A_1, E_3) + \frac{1}{2}\text{cov}(A_2, E_3)$. As the two parents play symmetric roles, $\text{cov}(A_1, E_3) = \text{cov}(A_2, E_3)$ and we get

$$\text{cov}(A_1, E_3) = \text{cov}(A_3, E_3). \tag{27}$$

We have $\text{cov}(A_3, E_3) = \rho(t+1)a(t+1)e$ (cf equation 22), thus

$$r_{A_1 E_3}(t) = \rho(t+1)\frac{a(t+1)}{a(t)}. \tag{28}$$

##### 3.6 Equation of evolution of the gene-environment correlation $\rho(t)$

Conditionnaly to $(E_1, E_2)$, $A_3$ and $E_3$ are independent. We are going to apply lemma 2 (appendix A), with $X = (A_1, E_3)$ and $Y = \frac{1}{e}(E_1, E_2)$. One has

$$\text{var}(Y) = \begin{pmatrix} 1 & r_{E_1E_2}(t) \\ r_{E_1E_2}(t) & 1 \end{pmatrix},$$

and

$$\begin{aligned}\text{cov}(X, Y) &= \frac{1}{e}\begin{pmatrix} \text{cov}(A_1, E_1) & \text{cov}(A_1, E_2) \\ \text{cov}(E_3, E_1) & \text{cov}(E_3, E_2) \end{pmatrix} \\ &= \begin{pmatrix} \rho(t)a(t) & r_{A_1E_2}(t)a(t) \\ ve & ve \end{pmatrix}.\end{aligned}$$

From lemma 2, we get

$$\begin{aligned}\text{cov}(A_1, E_3) &= \frac{\rho(t)a(t) + r_{A_1E_2}(t)a(t)}{1 + r_{E_1E_2}(t)}ve \\ &= \frac{\big(\rho(t) + r_{A_1E_2}(t)\big)a(t)}{1 + r_{E_1E_2}(t)}ve.\end{aligned}$$

We have $\text{cov}(A_1, E_3) = \text{cov}(A_3, E_3) = \rho(t+1)a(t+1)e$ (equations 27 and 22), thus we obtain

$$\rho(t+1) = v\frac{a(t)}{a(t+1)}\frac{\big(\rho(t) + r_{A_1E_2}(t)\big)}{1 + r_{E_1E_2}(t)},$$

or

$$\rho(t+1) = v\frac{a(t)}{a(t+1)}\frac{\rho(t) + \frac{r_{\text{ho}}}{\sigma^2(t)}\big(\rho(t)a(t)+e\big)\big(\rho(t)e + a(t)\big)}{1 + \frac{r_{\text{ho}}}{\sigma^2(t)}\big(\rho(t)a(t)+e\big)^2}, \tag{29}$$

where the value of $r_{A_1E_2}$ and $r_{E_1E_2}$ are taken from table 1 (and equations 19 and 18).

##### 3.7 Values at equilibrium

If one assumes there is equilibrium, that is all values don't depend on $t$ anymore, we can use equations 23, 24 and 28 to obtain some values at equilibrium.

From equation 23 we get

$$r_{\text{ga}} = \frac{1}{2} r_{A_1A_2}(1 + r_{\text{ga}}),$$

thus

$$r_{\text{ga}} = \frac{r_{A_1A_2}}{2 - r_{A_1A_2}} \tag{30}$$

or equivalently

$$r_{A_1A_2} = 2\frac{r_{\text{ga}}}{1 + r_{\text{ga}}}. \tag{31}$$

This gives a link between $r_{\text{ga}}$ and other model parameters, as we know that $r_{A_1A_2} = \frac{r_{\text{ho}}}{\sigma^2}(a + \rho e)^2$ (cf table 1), so for example

$$2\frac{r_{\text{ga}}}{1 + r_{\text{ga}}} = \frac{r_{\text{ho}}}{\sigma^2}(a + \rho e)^2$$

and

$$2r_{\text{ga}}(a^2 + 2\rho ae + e^2) = (1 + r_{\text{ga}})r_{\text{ho}}(a + \rho e)^2. \tag{32}$$

From equation 24, we get

$$r_{A_1A_3} = \frac{1}{2}(1 + r_{A_1A_2}) = \frac{1}{2}\left(1 + \frac{r_{\text{ho}}}{\sigma^2}(a + \rho e)^2\right).$$

From equation 28, we get $r_{A_1E_3} = \rho$. Note that the equilibrium hypothesis implies that we also have $r_{A_3E_3} = \rho$.

We have proved all the results summarized in table 2. We will show in sections 4 and 5 how to compute the equilibrium value of $\rho$.

### 4 Equilibrium values: equal variance case

In this section, we assume for the sake of simplicity that all loci effects have equal variance, that is, going back to the notations of sections 1.2 and 1.3, $\beta_i^2 p_i q_i$ doesn't depend on $i$, and thus for all $i$, $\kappa_i^2 = 1$. We will drop this assumption in section 5.

Note that the "natural" parameters of the model are $g_0$ (gametic standard deviation when there is no homogamy), $e$ (environmental standard deviation), $r_{\text{ho}}$ (correlation between parental phenotypes) and $v$ (environmental correlation between parent and offspring). These parameters don't depend on the generation $t$. Our aim here is to show that the equilibrium values of all other parameters can be computed from these four parameters.

We have derived evolution equations 21, 23, and 29, linking $r_{\text{ga}}(t)$, $a(t)$, and $\rho(t)$, but we have not yet a way to derive their equilibrium values. As we have $a^2(t) = 2(1 + r_{\text{ga}}(t))g^2(t)$, we will also consider here $g^2(t)$, the gametic variance.

The gametic variance $g^2(t)$ depends interloci on correlations, thus of the $\kappa_{ij}(t)$ introduced in section 1.3. As we assume here that for all $i$, $\kappa_i = 1$, from equation 5, we have

$$\mathrm{cor}\big(g_i^p, g_j^p\big) = \kappa_{ij}(t).$$

We will first establish in section 4.1 evolution equations for the $\kappa_{ij}(t)$.

We will establish in section 4.2 a link between the equilibrium value of the $\kappa_{ij}$ and the equilibrium value of the gametic correlation $r_{\mathrm{ga}}$. From this we will be able (section 4.3) to prove that the equilibrium values of $g^2(t)$ and $r_{\mathrm{ga}}(t)$ satisfy

$$g^2 = \frac{g_0^2}{1 - r_{\mathrm{ga}}}. \tag{33}$$

The equilibrium value of the genetic variance will then satisfy

$$a^2 = (1 + r_{\mathrm{ga}})2g^2 = \frac{1 + r_{\mathrm{ga}}}{1 - r_{\mathrm{ga}}} 2g_0^2. \tag{34}$$

Plugging in this equation what we have established in the previous section about the equilibrium value of $r_{\mathrm{ga}}$ (equations 30 and 32) leads to a (non-linear) system of equations in $a$ and $\rho$, which can be solved numerically to get all equilibrium values (section 4.4).

##### 4.1 Equation of evolution of inter-loci correlations $\kappa_{ij}(t)$

We consider here the inter-loci correlation. Note again that in the whole section, we assume that for all $i$, $\kappa_i = 1$.

It's the evolution of $\kappa_{ij}(t)$ that will be considered here: assuming a correlation $r_{\mathrm{ga}}(t) = \mathrm{cor}\big(G^p, G^m\big)$ between the genetic values of the parental gametes carried by an individual at the generation $t$, we will obtain a value for $\kappa_{ij}(t+1)$.

We need first to compute the correlation between loci on paternal and maternal gametes. We will then be able to use it to establish the equation of evolution.

###### 4.1.1 Correlation between loci on paternal and maternal gametes

As noted previously, we have $\mathrm{cor}\big(g_i^p, g_j^p\big) = \kappa_{ij}(t)$. As all loci play symmetric roles, and from

$$\mathrm{cov}\big(G^p, G^m\big) = \sum_{i,j} \mathrm{cov}\big(g_i^p, g_i^m\big) = r_{\mathrm{ga}}(t) g(t)^2,$$

we deduce

$$\mathrm{cov}\big(g_i^p, g_j^m\big) = \frac{1}{N^2} r_{\mathrm{ga}}(t) g(t)^2,$$

and as $g^2(t) = N g_0^2 \overline{\kappa}(t)$ (equation 6), we get for all $i$, $j$

$$\mathrm{cov}\big(g_i^p, g_j^m\big) = \frac{1}{N} g_0^2 r_{\mathrm{ga}}(t) \overline{\kappa}(t).$$

Dividing both size by $\mathrm{var}\big(g_i^p\big) = \mathrm{var}\big(g_j^m\big) = \frac{1}{N} g_0^2$ (equation 3) gives

$$\mathrm{cor}\big(g_i^p, g_j^m\big) = r_{\mathrm{ga}}(t) \overline{\kappa}(t). \tag{35}$$

##### 4.1.2 The equation of evolution of $\kappa_{ij}(t)$

Now we turn to the correlation between loci carried by the same gamete. Consider a gamete emitted an individual at generation $t$. Let $I_p$ and $I_m$ be as in subsection 1.6, and let the genetic value of the emitted gamete be $G = \sum_{i=1}^{N} g_i$ where $g_i = g_i^p$ if $i \in I_p$ and $g_i = g_i^m$ if $i \in I_m$. Then

- If $i, j \in I_p$, then $g_i = g_i^p$ et $g_j = g_j^p$ and $\text{cov}(g_i, g_j) = \text{cov}\left(g_i^p, g_j^p\right)$, thus $\kappa_{ij}(t+1) = \kappa_{ij}(t)$.
- The same holds if $i, j \in I_m$.
- If there is a recombination between paternal and maternal loci, that is if $i \in I_p, j \in I_m$ or $i \in I_m, j \in I_p$, we have

  $$\text{cov}(g_i, g_j) = \text{cov}\left(g_i^p, g_j^m\right) = \text{cov}\left(g_i^m, g_j^p\right),$$

  and, thank to equation 35, $\kappa_{ij}(t+1) = r_{\text{ga}}(t)\overline{\kappa}(t)$.

Finally, we have

$$\kappa_{ij}(t+1) = (1-\theta_{ij})\kappa_{ij}(t) + \theta_{ij} r_{\text{ga}}(t)\overline{\kappa}(t). \tag{36}$$

where $\theta_{ij}$ is the recombination rate between loci $i$ and $j$, which is the probability that loci $i, j$ are both in $I_p$ or both in $I_m$. We assume classically $0 < \theta_{ij} < \frac{1}{2}$ for $i \neq j$ and $\theta_{ii} = 0$. Note that for $i = j$ the above equation specializes into $\kappa_{ii}(t+1) = \kappa_{ii}(t) = 1$, which corresponds to the fact that allele frequencies don't change over time.

#### 4.2 Mean inter-loci correlation $\overline{\kappa}$ at equilibrium

Assuming that the population is at equilibrium, with $r_{\text{ga}}(t) = r_{\text{ga}}$, and $\overline{\kappa}(t) = \overline{\kappa}$ we get from equation 36, for all $i \neq j$,

$$\kappa_{ij} = (1-\theta_{ij})\kappa_{ij} + \theta_{ij} r_{\text{ga}}\overline{\kappa},$$

that is

$$\theta_{ji}\kappa_{ij} = \theta_{ij} r_{\text{ga}}\overline{\kappa},$$

and finally

$$\kappa_{ij} = r_{\text{ga}}\overline{\kappa}. \tag{37}$$

As $\kappa_{ii} = \kappa_i^2 = 1$ for all $i$, we get $\overline{\kappa} = \frac{1}{N} + \frac{N-1}{N} r_{\text{ga}}\overline{\kappa}$, and finally

$$\overline{\kappa} = \frac{1}{N-(N-1)r_{\text{ga}}} \simeq \frac{1}{N}\frac{1}{1-r_{\text{ga}}}. \tag{38}$$

The final approximation holds for large $N$, unless $r_{\text{ga}}$ is very close to 1. Namely, it holds if $N \gg (1-r_{\text{ga}})^{-1}$, which is a reasonnable assumption (see concluding remarks in section 4.3).

From equations 37 and 38 we get that the inter-loci correlations $\kappa_{ij}$ at equilibrium are all equal to $\frac{1}{N}\frac{r_{\text{ga}}}{1-r_{\text{ga}}}$, which is infinitesimal (of order of magnitude $\frac{1}{N}$), as expected.

Note that the correlation between loci on different gametes is also $r_{\text{ga}}\overline{\kappa}$, that is

$$\text{cov}\left(g_i^p, g_j^m\right) = \frac{1}{N}\frac{r_{\text{ga}}}{1-r_{\text{ga}}}.$$

This holds even for $i = j$, and shows that in this model, the departure from Hardy-Weinberg equilibrium induced by homogamy at a given loci is infinitesimal as well.

#### 4.3 Gametic variance at equilibrium

The gametic variance at equilibrium is then (equation 6 and 38)

$$g^2 = Ng_0^2\overline{\kappa} = \frac{g_0^2}{1-r_{\text{ga}}}. \tag{39}$$

The presence of a correlation between parental gametes increases the gametic variance, arising from the cumulative effect of infinitesimal correlations between all the causal loci. Moreover, one has $a^2 = 2(1+r_{\text{ga}})g^2$ (equation 7), and finally

$$a^2 = \frac{1+r_{\text{ga}}}{1-r_{\text{ga}}} \times 2g_0^2. \tag{40}$$

**Remarks.** Equation 30 gives $r_{\text{ga}} = \frac{r_{A_1A_2}}{2-r_{A_1A_2}}$, thus we get

$$a^2 = \frac{2g_0^2}{1-r_{A_1A_2}}. \tag{41}$$

This result is identical to what is derived, for example, in Crow and Felsenstein (1968) [1].

The approximation made above to express $\overline{\kappa}$ (equation 38), then $g^2$ and $a^2$, are valid unless $r_{\text{ga}}$ was very close to one. From equation 30, we see that this happens only in the marginal case were $r_{A_1A_2}$ is very close to one, which in turn happens only if $r_{\text{ho}}$ is very close to one and $e^2 \ll \sigma^2$ : the assortative mating is almost complete, and the environment has almost no part in the trait variance.

#### 4.4 Computing the equilibrium values of $a$ and $\rho$

We show here how the equilibrium values of $a$ and $\rho$ can be computed using the results already established. Recall that the parameters $g_0$, $r_{\text{ho}}$, $e$ and $v$ are given, and all other quantities in the model need to be derived from these.

##### 4.4.1 A system of two equations

We have shown in the above subsection (equations 40 and 41) that

$$a^2 = 2\frac{1+r_{\text{ga}}}{1-r_{\text{ga}}}g_0^2 = \frac{2g_0^2}{1-r_{A_1A_2}},$$

where $r_{A_1A_2}$ is (see table 1)

$$r_{A_1A_2} = \frac{r_{\text{ho}}}{\sigma^2}\left(a+\rho e\right)^2 = r_{\text{ho}}\frac{(a+\rho e)^2}{a^2+2\rho ae+e^2}.$$

Thus we have

$$a^2\left(1-r_{\text{ho}}\frac{(a+\rho e)^2}{a^2+2\rho ae+e^2}\right) = 2g_0^2. \tag{42}$$

Moreover, assuming equilibrium, equation 29 gives

$$\rho = v \frac{\rho + \frac{r_{ho}}{\sigma^2}(\rho a + e)(\rho e + a)}{1 + \frac{r_{ho}}{\sigma^2}(\rho a + e)^2},$$

which can be rewritten as

$$\rho \frac{\sigma^2 + r_{ho}(\rho a + e)^2}{\rho\sigma^2 + r_{ho}(\rho a + e)(\rho e + a)} = v \tag{43}$$

The two equations 42 and 43 give a system of two equations in $\rho$ and $a$.

##### 4.4.2 Solving the system numerically

We can first solve numerically equation 42 to get $a = a(\rho)$ as a function of $\rho$ (and of the known parameters $g_0^2$, $r_{ho}$ and $e$). This equation admits a unique solution $a(\rho) \in \left[0, \sqrt{\frac{2}{1-r_{ho}}} g_0\right]$.

Then, we plug this solution in equation 43. We obtain an equation where $\rho$ is the only unknown parameter:

$$\rho \frac{a(\rho)^2 + 2\rho a(\rho)e + e^2 + r\left(\rho a(\rho) + e\right)^2}{\rho\left(a(\rho)^2 + 2\rho a(\rho)e + e^2\right) + r\left(\rho a(\rho) + e\right)\left(\rho e + a(\rho)\right)} = v.$$

It can be numerically solved to get $\rho$ as a function of the known parameters $g_0^2$, $r_{ho}$, $e$ and $v$. Note that the left-hand side term varies from 0 for $\rho = 0$ to 1 for $\rho = 1$. Once the equation is solved for $\rho$, one naturally takes $a = a(\rho)$. All other parameters are then readily deduced from $a$ and $\rho$.

We provide R code for this resolution on the following github repository:

https://github.com/HervePerdry/AMVCTpaper

#### 4.5 Assortative mating without vertical cultural transmission

Here we look briefly at the particular case where $v = 0$. In that case, for all $t$ we have $\rho(t) = 0$, and many equations become simpler, allowing for explicit solutions.

In particular tables 1 and 2 can be summarized by the following table (the variable $E_3$ is not correlated to any of the other variables, and is not included).

| | $A_2$ | $A_3$ | $E_1$ | $E_2$ |
|---|---|---|---|---|
| $A_1$ | $r_{ho}\frac{a^2}{\sigma^2}$ | $\frac{1}{2}\left(1 + r_{ho}\frac{a^2}{\sigma^2}\right)$ | 0 | $r_{ho}\frac{ae}{\sigma^2}$ |
| $A_2$ | | $\frac{1}{2}\left(1 + r_{ho}\frac{a^2}{\sigma^2}\right)$ | $r_{ho}\frac{ae}{\sigma^2}$ | 0 |
| $A_3$ | | | $\frac{1}{2} r_{ho}\frac{ae}{\sigma^2}$ | $\frac{1}{2} r_{ho}\frac{ae}{\sigma^2}$ |
| $E_1$ | | | | $r_{ho}\frac{e^2}{\sigma^2}$ |

**Table 3:** Correlations at equilibrium between $A$ and $E$ terms in a trio without vertical cultural transmission

Equation 30 can be rewritten has

$$r_{\text{ga}} = \frac{r_{\text{ho}} \frac{a^2}{\sigma^2}}{2 - r_{\text{ho}} \frac{a^2}{\sigma^2}},$$

or equivalently

$$r_{\text{ga}} = \frac{r_{\text{ho}} a^2}{(2 - r_{\text{ho}}) a^2 + 2e^2}.$$

Moreover equation 42 reduces to

$$(1 - r_{\text{ho}}) a^4 + (e^2 - 2g_0^2) a^2 - 2g_0^2 = 0, \tag{44}$$

which is quadratic in $a^2$ and has only one positive solution:

$$a^2 = \frac{1}{2} \frac{1}{1 - r_{\text{ho}}} \left( \sqrt{(e^2 + 2g_0^2)^2 - 8 r_{\text{ho}} e^2 g_0^2} - (e^2 - 2g_0^2) \right). \tag{45}$$

#### 5 Equilibrium values: general case

We have proved in the previous section that when all loci have equal variance, the equilibrium value of $\overline{\kappa}$ is $(N(1 - r_{\text{ga}}))^{-1}$ (equation 38).

In sections 4.3 and 4.4, we used this result to show that $g^2 = \frac{g_0^2}{1 - r_{\text{ga}}}$ (equation 39), which in turn allowed us to derive a system of non linear equations for the equilibrium values of $\rho$ and $a$ (equations 42 and 43).

We show here that equation 38 (that is, $\overline{\kappa} = \frac{1}{N} \frac{1}{1 - r_{\text{ga}}}$) holds in the general case as well (equation 51), which will entail that the results of sections 4.3 and 4.4 still hold.

The following two sections are homologous of sections 4.1 and 4.2. and aim to prove that equation 38 hold in the general case.

We will also establish the equilibrium value of the inter-loci correlations $\text{cor}(g_i^p, g_j^p)$ (cf equation 53 below) and inter-gametic correlations $\text{cor}(g_i^p, g_j^m)$ (equation 54).

##### 5.1 Equation of evolution of inter-loci correlations $\kappa_{ij}(t)$

Here we no longer assume $\kappa_i = 1$ for all $i$, which gave us, by symmetry, $\text{cor}(g_i^p, g_j^m) = r_{\text{ga}} \overline{\kappa}$ (equation 35). In the general case, we have the more complicated value

$$\text{cov}(g_i^p, g_j^m) = \frac{1}{N} g_0^2 r_{\text{ga}} \frac{\overline{\kappa}_i \overline{\kappa}_j}{\overline{\kappa}}. \tag{46}$$

where $\overline{\kappa}_i = \frac{1}{N} \sum_{j=1}^N \kappa_{ij}$ and $\overline{\kappa} = \frac{1}{N^2} \sum_{i,j} \kappa_{ij}$. This result is proved in appendix A: see lemma 3, corollaries 1 and 2 which gives this result.

Following the same line of reasonning that in section 4.1.2, we get a new evolution equation:

$$\kappa_{ij}(t+1) = (1 - \theta_{ij}) \kappa_{ij}(t) + \theta_{ij} r_{\text{ga}}(t) \frac{\overline{\kappa}_i(t) \overline{\kappa}_j(t)}{\overline{\kappa}(t)}$$

where $\theta_{ij}$ is the recombination rate between loci $i$ and $j$. This gives

$$\kappa_{ij}(t+1) = \kappa_{ij}(t) + \theta_{ij} \left( r_{\text{ga}}(t) \frac{\overline{\kappa}_i(t) \overline{\kappa}_j(t)}{\overline{\kappa}(t)} - \kappa_{ij}(t) \right). \tag{47}$$

##### 5.2 Mean inter-loci correlation $\overline{\kappa}$ at equilibrium

The values at equilibrium can be obtained, assuming that the recombination rate $\theta_{ij}$ between loci $i$ and $j$ verifies $0 < \theta_{ij} < \frac{1}{2}$ for $i \neq j$. From equation 47 one gets that $\kappa_{ij}(t+1) = \kappa_{ij}(t) = \kappa_{ij}$ for $i \neq j$, if and only if

$$\kappa_{ij} = \kappa_{ij} + \theta_{ij}\left(r_{\text{ga}}\frac{\overline{\kappa}_i\overline{\kappa}_j}{\overline{\kappa}} - \kappa_{ij}\right)$$

that is

$$\kappa_{ij} = r_{\text{ga}}\frac{\overline{\kappa}_i\overline{\kappa}_j}{\overline{\kappa}} \ \text{ if } i \neq j$$

for $i \neq j$. On the other hand, $\kappa_{ii}(t) = \kappa_i^2$ does not depend on $t$, thus at equilibrium we still have $\kappa_{ii} = \kappa_i^2$, and finally

$$\kappa_{ij} = \begin{cases} r_{\text{ga}}\frac{\overline{\kappa}_i\overline{\kappa}_j}{\overline{\kappa}} & \text{if } i \neq j \\ \kappa_i^2 & \text{if } i = j \end{cases} \tag{48}$$

Summing this equation over $j$, we get

$$\sum_j \kappa_{ij} = \kappa_i^2 + r_{\text{ga}}\frac{\overline{\kappa}_i}{\overline{\kappa}}\sum_{j\neq i}\overline{\kappa}_j$$

Now the left-hand side of this equality is $N\overline{\kappa}_i$, and on the right-hand side the term $\sum_{j\neq i}\overline{\kappa}_j$ is equal to $N\overline{\kappa} - \overline{\kappa}_i$, hence

$$\overline{\kappa}_i = \frac{1}{N}\kappa_i^2 + r_{\text{ga}}\overline{\kappa}_i\left(1 - \frac{1}{N}\frac{\overline{\kappa}_i}{\overline{\kappa}}\right)$$

and

$$\left(\overline{\kappa}_i\right)^2 + N\overline{\kappa}\,\frac{1-r_{\text{ga}}}{r_{\text{ga}}}\overline{\kappa}_i - \frac{\overline{\kappa}}{r_{\text{ga}}}\kappa_i^2 = 0.$$

This second degree equation in $\overline{\kappa}_i$ has a single positive solution,

$$\overline{\kappa}_i = \sqrt{\left(\frac{1}{2}N\overline{\kappa}\frac{1-r_{\text{ga}}}{r_{\text{ga}}}\right)^2 + \frac{\overline{\kappa}}{r_{\text{ga}}}\kappa_i^2} - \frac{1}{2}N\overline{\kappa}\frac{1-r_{\text{ga}}}{r_{\text{ga}}} \tag{49}$$

Taking the mean of this over $i$, one gets an equation in $\overline{\kappa}$:

$$\overline{\kappa} = \frac{1}{N}\sum_{i=1}^{N}\sqrt{\left(\frac{1}{2}N\overline{\kappa}\frac{1-r_{\text{ga}}}{r_{\text{ga}}}\right)^2 + \frac{\overline{\kappa}}{r_{\text{ga}}}\kappa_i^2} - \frac{1}{2}N\overline{\kappa}\frac{1-r_{\text{ga}}}{r_{\text{ga}}}.$$

There is no closed form solution to this equation. However, it could be solved numerically to obtain the limiting value of $\overline{\kappa}$.

More simply, let $X = \frac{1}{2}N\overline{\kappa}\frac{1-r_{\text{ga}}}{r_{\text{ga}}}$ and $x = \frac{\overline{\kappa}}{r_{\text{ga}}}\kappa_i^2$, so that equation 49 becomes $\overline{\kappa}_i = \sqrt{X^2+x} - X$. We have $x \ll X$ as we assume $N$ is very large and $\kappa_i^2$ is close to 1. Using the first-order approximation $\sqrt{X^2+x} - X = \frac{x}{2X}$ for $x \ll X$ we get a simplified form of equation 49:

$$\overline{\kappa}_i \simeq \frac{\kappa_i^2}{N(1-r_{\text{ga}})} \tag{50}$$

and in turns, using $\frac{1}{N}\sum_i \kappa_i^2 = 1$, we have

$$\overline{\kappa} \simeq \frac{1}{N}\frac{1}{1-r_{\text{ga}}} \tag{51}$$

as in equation 38.

From equations 48, 50 and 51, we conclude that at equilibrium, for $i \neq j$,

$$\kappa_{ij} \simeq \frac{1}{N}\frac{r_{\text{ga}}}{1-r_{\text{ga}}}\kappa_i^2\kappa_j^2, \tag{52}$$

and finally, from equation 5, the inter-loci correlations are (for $i \neq j$)

$$\text{cor}\big(g_i^p, g_j^p\big) = \frac{\kappa_{ij}}{\kappa_i\kappa_j} \simeq \frac{1}{N}\frac{r_{\text{ga}}}{1-r_{\text{ga}}}\kappa_i\kappa_j. \tag{53}$$

Plugging equations 50 and 51 in equation 46, it comes that the correlation $\text{cor}\big(g_i^p, g_j^m\big)$ between loci on the paternal and maternal gametes is given by the same expression, even for $i = j$ :

$$\text{cor}\big(g_i^p, g_j^m\big) = \frac{\kappa_{ij}}{\kappa_i\kappa_j} \simeq \frac{1}{N}\frac{r_{\text{ga}}}{1-r_{\text{ga}}}\kappa_i\kappa_j. \tag{54}$$

### 6 Association statistics and polygenic scores

In this section (and the following), the population is assumed to be at equilibrium. In section 6.1 we are interested in the expected value of SNP heritability estimates. In section 6.2, we turn to the inflation of effect size estimates in association studies. In section 6.4, we are interested in the correlation between partial genetic values, for example between the odd and even numbered chromosomes.

#### 6.1 SNP-heritability

Consider pairs of unrelated individuals, with genetic correlation $\text{cor}(A_1, A_2) = u$. Despite the use of indices 1 and 2 as in sections 2 and 3, these individuals are not mates.

Haseman-Elston regression implies that the expected value of the SNP-heritability is

$$h^2_{SNP} = \frac{\text{cor}(P_1,P_2)}{\text{cor}(A_1,A_2)} = \frac{\text{cor}(P_1,P_2)}{u}.$$

We are going to compute $\text{cor}(P_1, P_2)$.

The individuals being unrelated, we assume that $E_1$ is independent of $E_2$ conditional to $A_2$. From lemma 1 it comes that

$$\text{cor}(E_1,E_2) = \text{cor}(E_1,A_2)\,\text{cor}(A_2,E_2).$$

We also have $A_1$ independent of $E_2$ conditional to $A_2$, and using again lemma 1,

$$\text{cor}(A_1,E_2) = \text{cor}(A_1,A_2)\,\text{cor}(A_2,E_2) = \rho u.$$

Similarly, we have $\text{cor}(A_2, E_1) = \rho u$. Finally

$$\text{cor}(E_1, E_2) = \text{cor}(E_1, A_2)\,\text{cor}(A_2, E_2) = \rho^2 u.$$

Then

$$\begin{aligned}\text{cov}(P_1, P_2) &= \text{cov}(A_1, A_2) + \text{cov}(A_1, E_2) + \text{cov}(E_2, A_1) + \text{cov}(E_1, E_2) \\ &= ua^2 + 2\rho uae + \rho^2 ue^2 = u(a + \rho e)^2.\end{aligned}$$

The expected value of the SNP-heritability is then

$$h^2_{SNP} = \frac{\text{cor}(P_1, P_2)}{\text{cor}(A_1, A_2)} = \frac{(a + \rho e)^2}{\sigma^2}. \tag{55}$$

Note that this quantity has been appearing previously in several places in the text, for example in equation 16. From equation 30, we get a link between the gametic correlation $r_{\text{ga}}$ and $h^2_{SNP}$:

$$r_{\text{ga}} = \frac{r_{\text{ho}} h^2_{SNP}}{2 - r_{\text{ho}} h^2_{SNP}}. \tag{56}$$

Similarly, from equation 41, we get that the genetic variance is

$$a^2 = \frac{2g_0^2}{1 - r_{\text{ho}} h^2_{SNP}}. \tag{57}$$

##### 6.2 Effect size estimates in association studies

We consider here the result of the regression of $P$ on the genotype $X_i = X_i^p + X_i^m$ at the $i$-th locus (cf section 1.2 for the notations) on $M$ unrelated individuals. The estimated effet $\hat{\beta}_i$ has expected value

$$E\left(\hat{\beta}_i\right) = \frac{\text{cov}(X_i, P)}{\text{var}(X_i)}.$$

We have $\text{cor}\left(X_i^p, X_i^m\right) = \text{cor}\left(g_i^p, g_i^m\right) = \frac{1}{N}\frac{r_{\text{ga}}}{1 - r_{\text{ga}}}\kappa_i\kappa_j \ll 1$ (equation 54). We have $\text{var}(X_i^p) = \text{var}(X_i^m) = p_i q_i$, thus the variance $\text{var}(X_i)$ is

$$\text{var}(X_i) = 2p_i q_i \left(1 + \frac{1}{N}\frac{r_{\text{ga}}}{1 - r_{\text{ga}}}\kappa_i\kappa_j\right) \simeq 2p_i q_i. \tag{58}$$

As mentioned before (section 4.2), the departure from Hardy-Weinberg equilibrium on a single locus is negligible.

Now turn to $\text{cov}(X_i, P)$. By symmetry, $X_i^p$ and $X_i^m$ playing the same rule, we have

$$\text{cov}(X_i, P) = \text{cov}\left(X_i^p + X_i^m, P\right) = 2\,\text{cov}\left(X_i^p, P\right). \tag{59}$$

Write $P = A + E = G^p + G^m + E$ and $\text{cov}\left(X_i^p, P\right) = \text{cov}\left(X_i^p, G^p\right) + \text{cov}\left(X_i^p, G^m\right) + \text{cov}\left(X_i^p, E\right)$. We need to compute three terms : $\text{cov}\left(X_i^p, G^p\right)$, $\text{cov}\left(X_i^p, G^m\right)$, and $\text{cov}\left(X_i^p, E\right)$.

##### 6.2.1 Computation of the first term: $\operatorname{cov}(X_i^p, G^p)$

As $g_i^p = \beta_i X_i^p$, we have, using equation 8

$$\operatorname{cov}(X_i^p, G^p) = \frac{1}{\beta_i}\operatorname{cov}(g_i^p, G^p)$$
$$= \frac{1}{\beta_i} g_0^2 \overline{\kappa}_i$$

and from equation 50

$$\operatorname{cov}(X_i^p, G^p) = \frac{1}{\beta_i} g_0^2 \frac{1}{N}\frac{\kappa_i^2}{1-r_{\text{ga}}}$$

but $\kappa_i^2 = N\frac{\beta_i^2 p_i q_i}{g_0^2}$ (equation 2), and we get

$$\operatorname{cov}(X_i^p, G^p) = \frac{\beta_i p_i q_i}{1-r_{\text{ga}}}. \tag{60}$$

##### 6.2.2 Computation of the second term: $\operatorname{cov}(X_i^p, G^m)$

Conditionnal to $G^p$, $X_i^p$ and $G^m$ are independent, thus (lemma 1)

$$\operatorname{cor}(X_i^p, G^m) = \operatorname{cor}(X_i^p, G^p)\operatorname{cor}(G^p, G^m) = r_{\text{ga}}\operatorname{cor}(X_i^p, G^p),$$

and, as $\operatorname{var}(G^p) = \operatorname{var}(G^m)$,

$$\operatorname{cov}(X_i^p, G^m) = r_{\text{ga}}\operatorname{cov}(X_i^p, G^p) \tag{61}$$

where $\operatorname{cov}(X_i^p, G^p)$ is given by equation 60.

##### 6.2.3 Computation of the third term: $\operatorname{cov}(X_i^p, E)$

Conditionnal to $A$, $X_i^p$ and $E$ are independent, thus (lemma 1),

$$\operatorname{cor}(X_i^p, E) = \operatorname{cor}(X_i^p, A)\operatorname{cor}(A, E) = \rho\operatorname{cor}(X_i^p, A),$$

and

$$\operatorname{cov}(X_i^p, E) = \rho\frac{e}{a}\operatorname{cov}(X_i^p, A).$$

Now thanks to equation 61

$$\operatorname{cov}(X_i^p, A) = \operatorname{cov}(X_i^p, G^p) + \operatorname{cov}(X_i^p, G^m) = (1 + r_{\text{ga}})\operatorname{cov}(X_i^p, G^p),$$

and

$$\operatorname{cov}(X_i^p, E) = \rho\frac{e}{a}(1 + r_{\text{ga}})\operatorname{cov}(X_i^p, G^p), \tag{62}$$

where $\operatorname{cov}(X_i^p, G^p)$ is given by equation 60.

##### 6.2.4 Conclusion of the computation: value of $\operatorname{cov}\left(X_i^p, P\right)$ and $E\left(\hat{\beta}_i\right)$

We have

$$\operatorname{cov}(X_i^p, P) = \operatorname{cov}\left(X_i^p, G^p\right) + \operatorname{cov}\left(X_i^p, G^m\right) + \operatorname{cov}\left(X_i^p, E\right)$$

thus using equations 61 and 62 we have

$$\operatorname{cov}(X_i^p, P) = \left(1 + r_{\mathrm{ga}} + \rho \frac{e}{a}(1 + r_{\mathrm{ga}})\right) \operatorname{cov}\left(X_i^p, G^p\right) \tag{63}$$

and using 60

$$\operatorname{cov}(X_i^p, P) = \frac{1 + r_{\mathrm{ga}}}{1 - r_{\mathrm{ga}}} \left(1 + \rho \frac{e}{a}\right) \beta_i p_i q_i. \tag{64}$$

From equations 58 and 59, we have

$$\begin{aligned} E\left(\hat{\beta}_i\right) &= \frac{\operatorname{cov}(X_i, P)}{\operatorname{var}(X_i)} \\ &= 2\frac{\operatorname{cov}\left(X_i^p, P\right)}{\operatorname{var}(X_i)} \\ &= \frac{1 + r_{\mathrm{ga}}}{1 - r_{\mathrm{ga}}} \left(1 + \rho \frac{e}{a}\right) \beta_i \end{aligned}$$

and finally

$$E\left(\hat{\beta}_i\right) = \gamma \beta_i \tag{65}$$

where we let the inflation factor $\gamma$ be

$$\gamma = \frac{1 + r_{\mathrm{ga}}}{1 - r_{\mathrm{ga}}} \left(1 + \rho \frac{e}{a}\right). \tag{66}$$

From equation 56, we get

$$\frac{1 + r_{\mathrm{ga}}}{1 - r_{\mathrm{ga}}} = \left(1 - r_{\mathrm{ho}} h^2_{SNP}\right)^{-1},$$

thus we can also write $\gamma$ as

$$\gamma = \frac{1 + \rho \frac{e}{a}}{1 - r_{\mathrm{ho}} h^2_{SNP}}. \tag{67}$$

#### 6.3 Polygenic scores

We consider a polygenic score is $\hat{A} = \sum_i X_i \hat{\beta}_i$ where the $\beta_i$ have been obtained on a sample of size $M$, independent of the sample on which the $\hat{A}$ is computed. We will compute here the (expected value of) the correlation $\operatorname{cor}\left(\hat{A}, P\right)$ between the polygenic score and the phenotype. In the absence of AM and VCT, our results are equivalent to those established established in Daetwyler et al (2008) [2].

For this, we first need to compute $\operatorname{var}\left(\hat{A}\right)$ and $\operatorname{cor}\left(\hat{A}, A\right)$.

##### 6.3.1 Variance $\text{var}(\hat{A})$ of the polygenic score

To compute the variance of $\hat{A}$, we will first compute the covariance $\text{var}(\hat{\beta}_i, \hat{\beta}_j)$ ; this will allow us to compute the variance of $\hat{A}$, conditional to the vector of genotypes ; then, by deconditionning, we will obtain $\text{var}(\hat{A})$.

We have (equation 65) $E(\hat{\beta}_i) = \gamma\beta_i$ and, classically, $\text{var}(\hat{\beta}_i) = \frac{\sigma^2}{M \cdot 2p_i q_i}$. The correlation $\text{cor}(\hat{\beta}_i, \hat{\beta}_j)$ between the SNP effect estimates for two different SNPs is approximately the linkage disequilibrium between loci $i$ and $j$:

$$\text{cor}(\hat{\beta}_i, \hat{\beta}_j) = \text{cor}(X_i, X_j) \tag{68}$$

Now

$$\begin{aligned}\text{cov}(X_i, X_j) &= \text{cov}(X_i^p + X_i^m, X_j^p + X_j^m) \\ &= \text{cov}(X_i^p, X_j^p) + \text{cov}(X_i^p, X_j^m) + \text{cov}(X_i^m, X_j^p) + \text{cov}(X_i^m, X_j^m).\end{aligned}$$

Thanks to equations 53 and 54, all these four terms can be computed, and all have the same value, for example

$$\begin{aligned}\text{cov}(X_i^p, X_j^p) &= \sqrt{p_i q_i \cdot p_j q_j}\,\text{cor}(X_i^p, X_j^p) \\ &= \sqrt{p_i q_i \cdot p_j q_j}\,\text{cor}(g_i^p, g_j^p) \\ &= \sqrt{p_i q_i \cdot p_j q_j}\,\frac{1}{N}\frac{r_{\text{ga}}}{1 - r_{\text{ga}}}\kappa_i\kappa_j\end{aligned}$$

and

$$\text{cov}(X_i, X_j) = 4\sqrt{p_i q_i \cdot p_j q_j}\,\frac{1}{N}\frac{r_{\text{ga}}}{1 - r_{\text{ga}}}\kappa_i\kappa_j. \tag{69}$$

As $\text{var}(X_i) = 2p_i q_i$ and $\text{var}(X_j) = 2p_j q_j$, we have

$$\text{cor}(X_i, X_j) = \frac{2}{N}\frac{r_{\text{ga}}}{1 - r_{\text{ga}}}\kappa_i\kappa_j.$$

Now we compute from the previous equation and from equation 68,

$$\text{cov}(\hat{\beta}_i, \hat{\beta}_j) = \frac{2}{N}\frac{r_{\text{ga}}}{1 - r_{\text{ga}}}\kappa_i\kappa_j\sqrt{\frac{\sigma^2}{M \cdot 2p_i q_i}}\sqrt{\frac{\sigma^2}{M \cdot 2p_j q_j}},$$

and

$$\text{cov}(\hat{\beta}_i, \hat{\beta}_j) = \frac{1}{N}\frac{r_{\text{ga}}}{1 - r_{\text{ga}}}\frac{\kappa_i\kappa_j}{\sqrt{p_i q_i \cdot p_j q_j}}\frac{\sigma^2}{M}. \tag{70}$$

Conditionnal to the vector of genotypes $\mathbf{X} = (X_1, \ldots, X_N)$, $A$ is a constant. The expected value of $\hat{A}$ conditional to $\mathbf{X}$ is

$$E(\hat{A} \mid \mathbf{X}) = \sum_i X_i E(\hat{\beta}_i) = \sum_i X_i \gamma\beta_i = \gamma A,$$

(equations 65 and 66), and its variance is

$$\begin{aligned}\text{var}(\hat{A} \mid \mathbf{X}) &= \sum_{i,j} X_i X_j \,\text{cov}(\hat{\beta}_i, \hat{\beta}_j) \\ &= \frac{\sigma^2}{M}\sum_i \frac{X_i^2}{2p_i q_i} + \frac{\sigma^2}{M}\frac{1}{N}\frac{r_{\text{ga}}}{1 - r_{\text{ga}}}\sum_{i \neq j} X_i X_j \frac{\kappa_i\kappa_j}{\sqrt{p_i q_i \cdot p_j q_j}}\end{aligned}$$

By deconditionning we get $E\left(\hat{A}\right) = E(E(A \mid X)) = \gamma E(A) = 0$, and

$$\begin{aligned}\operatorname{var}\left(\hat{A}\right) &= \operatorname{var}(E(A \mid X)) + E(\operatorname{var}(A \mid X))\\ &= \gamma^2 \operatorname{var}(A) + \frac{\sigma^2}{M}\left(\sum_i \frac{E(X_i^2)}{2p_i q_i} + \frac{1}{N}\frac{r_{\text{ga}}}{1-r_{\text{ga}}}\sum_{i\neq j} E(X_i X_j)\frac{\kappa_i \kappa_j}{\sqrt{p_i q_i \cdot p_j q_j}}\right)\end{aligned}$$

where $E(X_i^2) = 2p_i q_i$, and using equation 69 for $E(X_i X_j) = \operatorname{cov}(X_i, X_j)$, we have

$$E(X_i X_j)\frac{\kappa_i \kappa_j}{\sqrt{p_i q_i \cdot p_j q_j}} = \frac{4}{N}\frac{r_{\text{ga}}}{1-r_{\text{ga}}}\kappa_i^2 \kappa_j^2;$$

finally, we have

$$\operatorname{var}\left(\hat{A}\right) \simeq \gamma^2 a^2 + \frac{\sigma^2}{M}\left(N + \left(\frac{r_{\text{ga}}}{1-r_{\text{ga}}}\right)^2 \frac{4}{N^2}\sum_{i\neq j}\kappa_i^2 \kappa_j^2\right)$$

Now all $\kappa_i^2$ are close to 1, and $N$ is assumed to be very large, from which

$$\begin{aligned}\frac{4}{N^2}\sum_{i\neq j}\kappa_i^2 \kappa_j^2 &= \frac{2}{N^2}\left(\left(\sum_i \kappa_i^2\right)^2 - \sum_i \kappa_i^4\right)\\ &\simeq \frac{2}{N^2}\left(N^2 - N\right) \simeq 2.\end{aligned}$$

Thus

$$\operatorname{var}\left(\hat{A}\right) = \gamma^2 a^2 + \frac{\sigma^2}{M}\left(N + 2\left(\frac{r_{\text{ga}}}{1-r_{\text{ga}}}\right)^2\right)$$

and as $\frac{r_{\text{ga}}}{1-r_{\text{ga}}} \ll N$ (cf final comments of section 4.3) we have finally

$$\operatorname{var}\left(\hat{A}\right) \simeq \gamma^2 a^2 + \frac{N}{M}\sigma^2. \tag{71}$$

###### 6.3.2 Correlation $\operatorname{cor}\left(\hat{A}, A\right)$ between the polygenic score and the genetic value

We have $E(A \mid \mathbf{X}) = 0$, $E(\hat{A} \mid \mathbf{X}) = 0$, and $E(A\hat{A} \mid \mathbf{X}) = AE(\hat{A} \mid X) = \gamma A^2$. The law of total covariance gives $\operatorname{cov}\left(A, \hat{A}\right) = E(\gamma A^2) + \operatorname{cov}(0,0)$, and

$$\operatorname{cov}\left(\hat{A}, A\right) = \gamma a^2. \tag{72}$$

We get, using $\operatorname{var}(A) = a^2$ and $\operatorname{var}\left(\hat{A}\right)$ from equation 71,

$$\operatorname{cor}\left(\hat{A}, A\right) = \frac{\gamma a^2}{\sqrt{a^2\left(\gamma^2 a^2 + \frac{N}{M}\sigma^2\right)}} = \frac{\gamma a}{\sqrt{\gamma^2 a^2 + \frac{N}{M}\sigma^2}}. \tag{73}$$

We can also write it as

$$\operatorname{cor}\left(\hat{A}, A\right) = \frac{a}{\sqrt{a^2 + \frac{N}{M\gamma^2}\sigma^2}}. \tag{74}$$

It is interesting to replace $\gamma$ by its value (equation 67). We first write

$$\operatorname{cor}\left(\hat{A}, A\right)^2 = \frac{\gamma^2 a^2/\sigma^2}{\gamma^2 a^2/\sigma^2 + \frac{N}{M}}.$$

but

$$\begin{aligned}\frac{\gamma^2 a^2}{\sigma^2} &= \frac{\frac{a^2}{\sigma^2}\left(1+\rho\frac{e}{a}\right)^2}{\left(1-r_{\text{ho}}h^2_{SNP}\right)^2} \\ &= \frac{\frac{1}{\sigma^2}(a+\rho e)^2}{\left(1-r_{\text{ho}}h^2_{SNP}\right)^2},\end{aligned}$$

and from equation 55 we have

$$\frac{\gamma^2 a^2}{\sigma^2} = \frac{h^2_{SNP}}{\left(1-r_{\text{ho}}h^2_{SNP}\right)^2} \tag{75}$$

and finally

$$\operatorname{cor}\left(\hat{A}, A\right)^2 = \frac{h^2_{SNP}}{h^2_{SNP} + \frac{N}{M}\left(1-r_{\text{ho}}h^2_{SNP}\right)^2}. \tag{76}$$

##### 6.3.3 Correlation $\operatorname{cor}\left(\hat{A}, P\right)$ between the polygenic score and the phenotype

We have by lemma 1

$$\begin{aligned}\operatorname{cor}\left(\hat{A}, E\right) &= \operatorname{cor}\left(\hat{A}, A\right)\operatorname{cor}(A, E) \\ &= \rho \operatorname{cor}\left(\hat{A}, A\right) \\ &= \frac{\rho\gamma a}{\sqrt{\gamma^2 a^2 + \frac{N}{M}\sigma^2}}\end{aligned}$$

from equation 73. Thus, using equation 71,

$$\begin{aligned}\operatorname{cov}\left(\hat{A}, E\right) &= \operatorname{cor}\left(\hat{A}, E\right)\cdot\sqrt{\operatorname{var}\left(\hat{A}\right)}\cdot e \\ &= \frac{\rho\gamma a}{\sqrt{\gamma^2 a^2 + \frac{N}{M}\sigma^2}}\cdot\sqrt{\gamma^2 a^2 + \frac{N}{M}\sigma^2}\cdot e \\ &= \rho\gamma a e\end{aligned}$$

Using equation 72, we have

$$\begin{aligned}\operatorname{cov}\left(\hat{A}, P\right) &= \operatorname{cov}\left(\hat{A}, A\right) + \operatorname{cov}\left(\hat{A}, E\right) \\ &= \gamma a^2 + \rho e \gamma a\end{aligned}$$

that is

$$\operatorname{cov}\left(\hat{A}, P\right) = \gamma a(a+\rho e) \tag{77}$$

and, using equation 71,

$$\operatorname{cor}\left(\hat{A}, P\right) = \frac{\gamma a(a+\rho e)}{\sqrt{\gamma^2 a^2 + \frac{N}{M}\sigma^2}\cdot\sigma}. \tag{78}$$

Taking the square of this we get

$$
\begin{aligned}
\operatorname{cor}\left(\hat{A}, P\right)^2 &= \frac{\gamma^2 a^2}{\gamma^2 a^2 + \frac{N}{M}\sigma^2}\left(\frac{a+\rho e}{\sigma}\right)^2 \\
&= \frac{\gamma^2 a^2/\sigma^2}{\gamma^2 a^2/\sigma^2 + \frac{N}{M}} h^2_{SNP}
\end{aligned}
$$

and using equation 75

$$
\operatorname{cor}\left(\hat{A}, P\right)^2 = \frac{h^2_{SNP}}{h^2_{SNP} + \frac{N}{M}\left(1 - r_{\text{ho}} h^2_{SNP}\right)^2} h^2_{SNP}. \tag{79}
$$

#### 6.4 Partial genetic values and partial genetic scores

Here we consider to disjoints subsets of causal variants, $S_1, S_2 \subset \{1, \ldots, N\}$. In practice $S_1$ and $S_2$ could be the variants located on odd and even chromosomes, respectively.

We assume each $S_i$ contains a proportion $\alpha_i = \frac{1}{N}|S_i|$ of the causal variants. For $S = S_1$ or $S_2$, we denote

$$
G_S^p = \sum_{i\in S} g_i^p,\ G_S^m = \sum_{i\in S} g_i^m,\ \text{and } A_S = G_S^p + G_S^m.
$$

the partial genetic values associated to the subset $S$.

Our aim is to compute the correlation between $A_{S_1}$ and $A_{S_2}$. For this, we will first compute the variance of a parental gamete ($G_S^p$ or $G_S^m$), from which well deduce the variance of $A_S$, for $S = S_1$ or $S_2$. We will then be able to complete our computation.

Finally, we will compute the correlation between the partial genetic scores $\hat{A}_{S_1}$ and $\hat{A}_{S_2}$.

In the particular case where there is AM and no VCT, our results are similar to those of Yengo et al (2018) [5].

##### 6.4.1 Variance of the partial genetic value of a parental gamete

Let $S$ be a subset of causal variants. We have naturally $\operatorname{var}(G_S^p) = \operatorname{var}(G_S^m)$. We will compute for example $G_S^p$.

Using equations 3, 4, and 52, we have

$$
\begin{aligned}
\operatorname{var}(G_S^p) &= \sum_{i\in S}\operatorname{var}(g_i^p) + \sum_{i\neq j\in S}\operatorname{cov}(g_i^p, g_j^p) \\
&= \frac{1}{N} g_0^2\left(\sum_{i\in S}\kappa_i^2 + \sum_{i\neq j\in S}\kappa_{ij}\right) \\
&= \frac{1}{N} g_0^2\left(\sum_{i\in S}\kappa_i^2 + \frac{1}{N}\frac{r_{\text{ga}}}{1-r_{\text{ga}}}\sum_{i\neq j\in S}\kappa_i^2\kappa_j^2\right).
\end{aligned}
$$

As $S = S_1$ or $S_2$ contains a proportion $\alpha = \alpha_1$ or $\alpha_2$ of the causal variants, and $\sum_i \kappa_i^2 = N$ with all the $\kappa_i^2$ close to one, we have

$$
\sum_{i\in S}\kappa_i^2 \simeq N\alpha \ \text{ and } \sum_{i\neq j\in S}\kappa_i^2\kappa_j^2 \simeq \alpha^2 N^2, \tag{80}
$$

thus

$$\mathrm{var}\left(G_S^p\right) \simeq g_0^2\left(\alpha+\alpha^2\frac{r_{\mathrm{ga}}}{1-r_{\mathrm{ga}}}\right)$$

and finally

$$\mathrm{var}\left(G_S^p\right) \simeq \alpha g_0^2\left(1+\alpha\frac{r_{\mathrm{ga}}}{1-r_{\mathrm{ga}}}\right). \tag{81}$$

##### 6.4.2 Variance of the partial genetic value $A_S$

Using equations 3, 52, and 54,

$$\begin{aligned}\mathrm{cov}\left(G_S^p, G_S^m\right) &= \sum_{i,j\in S}\mathrm{cov}\left(g_i^p, g_j^m\right)\\ &= \frac{1}{N^2}g_0^2\frac{r_{\mathrm{ga}}}{1-r_{\mathrm{ga}}}\sum_{i,j\in S}\kappa_i^2\kappa_j^2\\ &\simeq \alpha^2 g_0^2\frac{r_{\mathrm{ga}}}{1-r_{\mathrm{ga}}},\end{aligned}$$

where we used equation 80 in the last step. Thus, using this and equation 81,

$$\begin{aligned}\mathrm{var}(A_S) &= \mathrm{var}\left(G_S^p\right)+2\,\mathrm{cov}\left(G_S^p, G_S^m\right)+\mathrm{var}\left(G_S^m\right)\\ &\simeq 2\alpha g_0^2\left(1+\alpha\frac{r_{\mathrm{ga}}}{1-r_{\mathrm{ga}}}\right)+2\alpha^2 g_0^2\frac{r_{\mathrm{ga}}}{1-r_{\mathrm{ga}}}\end{aligned}$$

and

$$\mathrm{var}(A_S) \simeq 2\alpha g_0^2\left(1+2\alpha\frac{r_{\mathrm{ga}}}{1-r_{\mathrm{ga}}}\right).$$

or equivalently

$$\mathrm{var}(A_S) \simeq 2\alpha g_0^2\left(\frac{1+(2\alpha-1)r_{\mathrm{ga}}}{1-r_{\mathrm{ga}}}\right). \tag{82}$$

Using equation 40, one can rewrite this as

$$\mathrm{var}(A_S) \simeq \alpha a^2\frac{1+(2\alpha-1)r_{\mathrm{ga}}}{1+r_{\mathrm{ga}}}. \tag{83}$$

##### 6.4.3 Correlation between the partial genetic values $A_{S_1}$ and $A_{S_2}$

As $A_{S_1} = G_{S_1}^p + G_{S_1}^m$ and $A_{S_2} = G_{S_2}^p + G_{S_2}^m$, we have

$$\mathrm{cov}(A_{S_1}, A_{S_2}) = \mathrm{cov}\left(G_{S_1}^p, G_{S_2}^p\right)+\mathrm{cov}\left(G_{S_1}^p, G_{S_2}^m\right)+\mathrm{cov}\left(G_{S_1}^m, G_{S_2}^p\right)+\mathrm{cov}\left(G_{S_1}^m, G_{S_2}^m\right) \tag{84}$$

We have $\mathrm{cov}\left(G_{S_1}^p, G_{S_2}^p\right) = \sum_{i\in S_1, j\in S_2}\mathrm{cov}\left(g_i^p, g_j^p\right)$, and using equation 4

$$\mathrm{cov}\left(G_{S_1}^p, G_{S_2}^p\right) = \frac{1}{N}g_0^2\sum_{i\in S_1, j\in S_2}\kappa_{ij}, \tag{85}$$

thus from equation 52,

$$\operatorname{cov}\left(G^p_{S_1}, G^p_{S_2}\right) = \frac{1}{N^2} g_0^2 \frac{r_{\text{ga}}}{1-r_{\text{ga}}} \sum_{i\in S_1, j\in S_2} \kappa_i^2 \kappa_j^2,$$

and using equation 80,

$$\operatorname{cov}\left(G^p_{S_1}, G^p_{S_2}\right) \simeq \alpha_1\alpha_2 g_0^2 \frac{r_{\text{ga}}}{1-r_{\text{ga}}}. \tag{86}$$

This is also the value of $\operatorname{cov}(G^m_{S_1}, G^m_{S_2})$.

But now thanks to equations 3 and 54 we have

$$\begin{aligned}\operatorname{cov}\left(G^p_{S_1}, G^m_{S_2}\right) &= \sum_{i\in S_1, j\in S_2} \operatorname{cov}\left(g^p_i, g^m_j\right)\\ &= \frac{1}{N} g_0^2 \sum_{i\in S_1, j\in S_2} \kappa_{ij},\end{aligned}$$

which is the same as equation 85; thus all four terms in equation 84 are equal, and we have

$$\operatorname{cov}(A_{S_1}, A_{S_2}) = 4\alpha_1\alpha_2 g_0^2 \frac{r_{\text{ga}}}{1-r_{\text{ga}}}. \tag{87}$$

Finally the correlation between the partial genetic values $A_{S_1}$ and $A_{S_2}$ is

$$\operatorname{cor}(A_{S_1}, A_{S_2}) = \frac{2\sqrt{\alpha_1\alpha_2}\, r_{\text{ga}}}{\sqrt{(1+(2\alpha_1-1)r_{\text{ga}})(1+(2\alpha_2-1)r_{\text{ga}})}}. \tag{88}$$

In the particular case where $\alpha_1 = \alpha_2 = \frac{1}{2}$, which corresponds approximately to splitting the autosome into odd and even chromosomes, this gives $\operatorname{cor}(A_{S_1}, A_{S_2}) = r_{\text{ga}}$.

##### 6.4.4 Correlation between the partial genetic scores $\hat{A}_{S_1}$ and $\hat{A}_{S_2}$

When using polygenic scores $\hat{A}_{S_1}$ and $\hat{A}_{S_2}$, this correlation has to be mitigated by factors corresponding to $\operatorname{cor}\left(\hat{A}_{S_1}, A_{S_1}\right)$, and $\operatorname{cor}\left(\hat{A}_{S_2}, A_{S_2}\right)$.

The computations of section 6.3 lead, *mutatis mutando,* to

$$\operatorname{var}\left(\hat{A}_S\right) = \gamma^2 \operatorname{var}(A_S) + \frac{N}{M}\alpha\sigma^2,$$

and

$$\operatorname{cov}\left(\hat{A}_S, A_S\right) = \gamma \operatorname{var}(A_S),$$

thus

$$\operatorname{cor}\left(\hat{A}_S, A_S\right)^2 = \frac{\gamma^2 \operatorname{var}(A_S)}{\gamma^2 \operatorname{var}(A_S) + \frac{N}{M}\alpha\sigma^2}.$$

By using equation 83 for $\operatorname{var}(A_S)$ one gets

$$\operatorname{cor}\left(\hat{A}_S, A_S\right)^2 = \frac{(1+(2\alpha-1)r_{\text{ga}})\gamma^2 a^2/\sigma^2}{(1+(2\alpha-1)r_{\text{ga}})\gamma^2 a^2/\sigma^2 + \frac{N}{M}(1+r_{\text{ga}})}.$$

Using equation 75 for $\gamma^2 a^2/\sigma^2$, we have

$$\operatorname{cor}\left(\hat{A}_S, A_S\right)^2 = \frac{(1+(2\alpha-1)r_{\text{ga}})h^2_{SNP}}{(1+(2\alpha-1)r_{\text{ga}})h^2_{SNP} + \frac{N}{M}(1+r_{\text{ga}})\left(1-r_{\text{ho}}h^2_{SNP}\right)^2}.$$

Now for the particular case where $\alpha = \frac{1}{2}$, we have

$$\operatorname{cor}\left(\hat{A}_S, A_S\right)^2 = \frac{h^2_{SNP}}{h^2_{SNP} + \frac{N}{M}(1+r_{\text{ga}})\left(1-r_{\text{ho}}h^2_{SNP}\right)^2},$$

and the correlation between partial genetic scores covering each half of the genome is

$$\operatorname{cor}\left(\hat{A}_{S_1}, \hat{A}_{S_2}\right) = \frac{h^2_{SNP}}{h^2_{SNP} + \frac{N}{M}(1+r_{\text{ga}})\left(1-r_{\text{ho}}h^2_{SNP}\right)^2} r_{\text{ga}}. \tag{89}$$

### 7 Within-family estimates

In this section again the population is assumed to be at equilibrium. In section 7.2 we consider the regression of the offspring phenotype on the mid-parent genotype; we show that while with AM only, it gives an unbiased estimate of heritability, this is no longer the case with VCT. In sections 7.3 and 7.4, we consider two types of family-based GWAS, in which an adjustment is done on parental and on offspring genotypes, respectively; and in 7.5, we consider the correlation between sibling difference in genotypes and sibling differences in genetic values.

#### 7.1 Notations

Contrarily to sections 2 and 3 where focus was on the couple, here focus is on the individual, the genotypes and phenotypes of their parents being used for adjustment. Consequently we will denote $A$ the genetic value of the individual, $E$ the contribution of their environment to their phenotype $P = A + E$; the corresponding quantities for their parents are denoted $A_p, A_m, G_p$, and so on.

As before, $A$ can be decomposed in $A = G^p + G^m$ where $G^p$ and $G^m$ are the genetic values of the parental gametes, and the parental genetic values can be written $A_p = G^p + H^p$, $A_m = G^m + H^m$, where $H^p$ and $H^m$ are the genetic value of the “untransmitted gametes" (cf section 1.6).

As in section 1.2, the individual genotype at a loci of index $i$ is $X_i$, with $X_i = X_i^p + X_i^m$ where $X_i^p$ and $X_i^m$ are the centered variable encoding the allele transmitted by the father and the mother, respectively; we denote by $Y_i^p$ and $Y_i^m$ the centered variable encoding the parental untransmitted alleles, so that the genotype of the two parents are $X_{pi} = X_i^p + Y_i^p$ and $X_{mi} = X_i^m + Y_i^m$.

We let (again as in section 1.2) $g_i^p = \beta_i X_i^p$, $g_i^m = \beta_i X_i^m$, and moreover we let $h_i^p = \beta_i Y_i^p$ and $h_i^m = \beta_i Y_i^m$, so that the genetic values of the untransmitted alleles are $H^p = \sum_i h_i^p$ and $H^m = \sum_i h_i^m$ (section 1.6).

#### 7.2 Regression on mid-parent phenotype

It is known that in presence of assortative mating, the coefficient of regression of the offspring phenotypes $P$ on the so-called mid-parent phenotype $P_{\text{mid}} = \frac{1}{2}(P_p + P_m)$ is an unbiased estimate of heritability when there is no shared environment.

The expected value $h^2_{MP}$ of this coefficient is

$$\begin{aligned} h^2_{MP} &= \frac{\text{cov}(P, P_{\text{mid}})}{\text{var}(P_{\text{mid}})} \\ &= 2\frac{\text{cov}(P, P_p) + \text{cov}(P, P_m)}{\text{var}(P_p + P_m)} \end{aligned}$$

We have $\text{cor}(P_p, P_m) = r_{\text{ho}}$ and $\text{var}(P_p) = \text{var}(P_m) = \sigma^2$, thus $\text{var}(P_p + P_m) = 2(1 + r_{\text{ho}})\sigma^2$, and by symmetry of the parents' roles, $\text{cov}(P, P_m) = \text{cov}(P, P_p)$, and

$$h^2_{MP} = 2\frac{\text{cov}(P, P_p)}{(1 + r_{\text{ho}})\sigma^2}.$$

We now compute $\text{cov}(P, P_p)$:

$$\begin{aligned} \text{cov}(P, P_p) &= cov(A + E, A_p + E_p) \\ &= r_{AA_p} a^2 + r_{AE_p} ae + r_{EA_p} ae + r_{EE_p} e^2 \\ &= \frac{1}{2}\left(1 + \frac{r_{\text{ho}}}{\sigma^2}(a + \rho e)^2\right) a^2 + \frac{1}{2}(\rho + \frac{r_{\text{ho}}}{\sigma^2}(a + \rho e)(\rho a + e))ae + \rho ae + \nu e^2 \\ &= \frac{1}{2}a^2 + \frac{1}{2}\frac{r_{\text{ho}}}{\sigma^2}(a + \rho e)a\left((a + \rho e)a + (\rho a + e)e\right) + \frac{3}{2}\rho ae + \nu e^2 \\ &= \frac{1}{2}a^2 + \frac{1}{2}\frac{r_{\text{ho}}}{\sigma^2}(a + \rho e)a\sigma^2 + \frac{3}{2}\rho ae + \nu e^2 \\ &= \frac{1}{2}a^2 + \frac{1}{2}r_{\text{ho}}(a + \rho e)a + \frac{3}{2}\rho ae + \nu e^2 \\ &= \frac{1}{2}(1 + r_{\text{ho}})a^2 + \frac{1}{2}(3 + r_{\text{ho}})\rho ae + \nu e^2 \end{aligned}$$

where the values of $r_{AA_p}$, $r_{AE_p}$ and $r_{EA_p}$ are taken from table 2, in which they are denoted $r_{A_3A_1}$, $r_{A_3E_1}$ and $r_{E_3A_1}$ respectively.

Thus we have

$$h^2_{MP} = \frac{(1 + r_{\text{ho}})a^2 + (3 + r_{\text{ho}})\rho ae + 2\nu e^2}{(1 + r_{\text{ho}})\sigma^2},$$

and finally

$$h^2_{MP} = h^2 + \frac{(3 + r_{\text{ho}})\rho ae + 2\nu e^2}{(1 + r_{\text{ho}})\sigma^2}. \tag{90}$$

In presence of AM without VCT, that is when $\nu$ is zero, and thus $\rho$ is zero too, we have $h^2_{MP} = h^2$, which is a classical result. We see here that this does not longer hold in presence of vertical cultural transmission (that is, if $\nu > 0$).

##### 7.3 Association study with adjustement on parental genotypes

Here we consider the regression of $P$ on the genotype $X_i$ of the variant of index $i$, and on $M_i = \frac{1}{2}(X_{pi} + X_{mi})$, the mean of the parental genotypes.

We will show that thanks to this "adjustment on $M_i$", the bias that we have brought to light in section 6.2 no longer exists. This result is similar to the result of Young et al (2022) [6].

Standard results on linear regression gives, for the regression of $P$ on two predictors denoted $\xi_1$ and $\xi_2$,

$$E\begin{pmatrix}\hat{\beta}_1\\ \hat{\beta}_2\end{pmatrix} = \begin{pmatrix}\operatorname{var}(\xi_1) & \operatorname{cov}(\xi_1,\xi_2)\\ \operatorname{cov}(\xi_1,\xi_2) & \operatorname{var}(\xi_2)\end{pmatrix}^{-1}\begin{pmatrix}\operatorname{cov}(\xi_1,P)\\ \operatorname{cov}(\xi_2,P)\end{pmatrix}. \tag{91}$$

We let here $\xi_1 = X_i$ and $\xi_2 = M_i$. We need first to compute the matrix

$$\begin{pmatrix}\operatorname{var}(X_i) & \operatorname{cov}(X_i,M_i)\\ \operatorname{cov}(X_i,M_i) & \operatorname{var}(M_i)\end{pmatrix}.$$

Using the notations defined in 7.1, we have

$$M_i = \frac{1}{2}\left(X_i^p + Y_i^p + X_i^m + Y_i^m\right).$$

We have showed in section 6.2 that the correlation between $X_i^p$ and $X_i^m$ can be neglected, so that $\operatorname{var}(X_i) = \operatorname{var}(X_i + X_i^m) \simeq p_i q_i$ (equation 58); similarly, the correlation between the four variables in $M_i$ can be neglected and we have $\operatorname{var}(M_i) \simeq p_i q_i$, and

$$\operatorname{cov}(X_i, M_i) \simeq \frac{1}{2}\left(\operatorname{var}\left(X_i^p\right) + \operatorname{var}\left(X_i^m\right)\right) = p_i q_i.$$

Thus we have

$$\begin{pmatrix}\operatorname{var}(X_i) & \operatorname{cov}(X_i,M_i)\\ \operatorname{cov}(X_i,M_i) & \operatorname{var}(Mi)\end{pmatrix} = p_i q_i \begin{pmatrix}2 & 1\\ 1 & 1\end{pmatrix}.$$

The inverse of this matrix is easy to compute; we get

$$E\begin{pmatrix}\hat{\beta}_i\\ \hat{\beta}'_i\end{pmatrix} = \frac{1}{p_i q_i}\begin{pmatrix}1 & -1\\ -1 & 2\end{pmatrix}\begin{pmatrix}\operatorname{cov}(X_i,P)\\ \operatorname{cov}(M_i,P)\end{pmatrix},$$

so

$$E\left(\hat{\beta}_i\right) = \frac{1}{p_i q_i}\left(\operatorname{cov}(X_i,P) - \operatorname{cov}(M_i,P)\right) = \frac{1}{p_i q_i}\operatorname{cov}(X_i - M_i, P),$$

and

$$E\left(\hat{\beta}'_i\right) = \frac{1}{p_i q_i}\left(-\operatorname{cov}(X_i,P) + 2\operatorname{cov}(M_i,P)\right) = \frac{1}{p_i q_i}\operatorname{cov}(2M_i - X_i, P).$$

We are primarily interested by showing that $E\left(\hat{\beta}_i\right) = \beta_i$.

We have

$$X_i - M_i = \frac{1}{2}\left(X_i^p - Y_i^p + X_i^m - Y_i^m\right),$$

and by symmetry of the two parents,

$$\operatorname{cov}\left(X_i^p - Y_i^p, P\right) = \operatorname{cov}\left(X_i^m - Y_i^m, P\right),$$

thus

$$E\left(\hat{\beta}_i\right) = \frac{1}{p_i q_i} \operatorname{cov}\left(X_i^p - Y_i^p, P\right). \tag{92}$$

We know (cf equation 63) that

$$\operatorname{cov}\left(X_i^p, P\right) = (1 + r_{\text{ga}}) \left(1 + \rho \frac{e}{a}\right) \operatorname{cov}\left(X_i^p, G^p\right)$$

where $\operatorname{cov}\left(X_i^p, G^p\right) = \frac{\beta_i p_i q_i}{1 - r_{\text{ga}}}$ (equation 60). Now we turn to $\operatorname{cov}\left(Y_i^p, P\right)$. Writing $P = G^p + G^m + E$, we have $\operatorname{cov}\left(Y_i^p, P\right) = \operatorname{cov}\left(Y_i^p, G^p\right) + \operatorname{cov}\left(Y_i^p, G^m\right) + \operatorname{cov}\left(Y_i^p, E\right)$. We need to compute three terms: $\operatorname{cov}\left(Y_i^p, G^p\right)$, $\operatorname{cov}\left(Y_i^p, G^m\right)$, and $\operatorname{cov}\left(Y_i^p, E\right)$.

###### 7.3.1 Computation of the first term: $\operatorname{cov}\left(Y_i^p, G^p\right)$

Conditionnal to $H^p$, $h_i^p$ and $G^p$ are independent, thus so are $Y_i^p$ and $G_p$. Thus by lemma 1,

$$\operatorname{cor}\left(Y_i^p, G^p\right) = \operatorname{cor}\left(Y_i^p, H^p\right) \operatorname{cor}\left(G^p, H^p\right),$$

We know that $\operatorname{cor}\left(G^p, H^p\right) = r_{\text{ga}}$ (equation 10). On the other hand, by symmetry, $\operatorname{cor}\left(Y_i^p, H^p\right) = \operatorname{cor}\left(X_i^p, G^p\right)$. So we have

$$\operatorname{cor}\left(Y_i^p, G^p\right) = r_{\text{ga}} \operatorname{cor}\left(X_i^p, G^p\right),$$

and

$$\operatorname{cov}\left(Y_i^p, G^p\right) = r_{\text{ga}} \operatorname{cov}\left(X_i^p, G^p\right). \tag{93}$$

###### 7.3.2 Computation of the second term: $\operatorname{cov}\left(Y_i^p, G^m\right)$

In the same way, $Y_i^p$ and $G^m$ are independent conditional to $H^p$, and we have

$$\operatorname{cor}\left(Y_i^p, G^m\right) = \operatorname{cor}\left(Y_i^p, H^p\right) \operatorname{cor}\left(G^m, H^p\right).$$

As before, $\operatorname{cor}\left(Y_i^p, H^p\right) = \operatorname{cor}\left(X_i^p, G^p\right)$, and the gametic values $G^p$ and $H^p$ play symmetric roles as far as $G^m$ is concerned thus $\operatorname{cor}\left(G^m, H^p\right) = \operatorname{cor}\left(G^m, G^p\right) = r_{\text{ga}}$ and

$$\operatorname{cov}\left(Y_i^p, G^m\right) = r_{\text{ga}} \operatorname{cov}\left(X_i^p, G^p\right). \tag{94}$$

###### 7.3.3 Computation of the third term: $\operatorname{cov}\left(Y_i^p, E\right)$

From the symmetric roles of $X_i^p$ and $Y_i^p$, we get that $\operatorname{cov}\left(Y_i^p, E\right) = \operatorname{cov}\left(X_i^p, E\right)$ (given by equation 62), so

$$\operatorname{cov}\left(Y_i^p, E\right) = (1 + r_{\text{ga}}) \rho \frac{e}{a} \operatorname{cov}\left(X_i^p, G^p\right). \tag{95}$$

###### 7.3.4 Conclusion of the computation: value of $\operatorname{cov}\left(Y_i^p, P\right)$ and $E\left(\hat{\beta}_i\right)$

Putting all three terms together, we have

$$\begin{aligned} \operatorname{cov}\left(Y_i^p, P\right) &= \operatorname{cov}\left(Y_i^p, G^p\right) + \operatorname{cov}\left(Y_i^p, G^m\right) + \operatorname{cov}\left(Y_i^p, E\right) \\ &= \left(2 r_{\text{ga}} + (1 + r_{\text{ga}}) \rho \frac{e}{a}\right) \operatorname{cov}\left(X_i^p, G^p\right) \end{aligned}$$

We have (equation 92)

$$E\left(\hat{\beta}_i\right) = \frac{1}{p_i q_i}\left(\text{cov}\left(X_i^p, P\right) - \text{cov}\left(Y_i^p, P\right)\right)$$

But using equation 63, we have

$$\begin{aligned}\text{cov}\left(X_i^p, P\right) - \text{cov}\left(Y_i^p, P\right) &= \left((1+r_{\text{ga}})\left(1+\rho\frac{e}{a}\right) - 2r_{\text{ga}} - (1+r_{\text{ga}})\rho\frac{e}{a}\right)\text{cov}\left(X_i^p, G^p\right) \\ &= (1-r_{\text{ga}})\,\text{cov}\left(X_i^p, G^p\right)\end{aligned}$$

and using equation 60,

$$\text{cov}\left(X_i^p, P\right) - \text{cov}\left(Y_i^p, P\right) = \beta_i p_i q_i \tag{96}$$

and finally get the desired result,

$$E\left(\hat{\beta}_i\right) = \beta_i.$$

##### 7.3.5 Coefficient of the adjustement variable $M_i$

As final note, the expected value of $\hat{\beta}'_i$, the coefficient of $M_i$, is

$$\begin{aligned}E\left(\hat{\beta}'_i\right) &= \frac{1}{p_i q_i}\text{cov}(2M_i - X_i, P) \\ &= \frac{1}{p_i q_i}\text{cov}(Y_i^p + Y_i^m, P) \\ &= \frac{2}{p_i q_i}\text{cov}(Y_i^p, P) \\ &= 2\frac{2r_{\text{ga}} + (1+r_{\text{ga}})\rho\frac{e}{a}}{1-r_{\text{ga}}}\beta_i \\ &= 2(\gamma - 1)\beta_i.\end{aligned}$$

where $\gamma = \frac{1+r_{\text{ga}}}{1-r_{\text{ga}}}\left(1+\rho\frac{e}{a}\right)$ is the inflation factor of the regression on $X$.

#### 7.4 Association study with adjustement on sibling genotypes

Instead on adjusting the mean parent genotype as in section 7.3 another possibility is to adjust on $S_i$, where $S_i$ is the mean of the genotypes of the individual and their $n-1 \geq 1$ sibs. We will show that in this way again, the bias in the estimation of the effect $\beta_i$ disappears. This result is similar to the result of Howe et al (2022) [3].

Let us denote by $Z_i^1, \ldots, Z_i^{n-1}$ the sibling genotypes, and let

$$S_i = \frac{1}{n}\left(X_i + \sum_{j=1}^{n-1} Z_i^j\right).$$

We will use equation 91, so we need to compute the matrix

$$\begin{pmatrix}\text{var}(X_i) & \text{cov}(X_i, S_i) \\ \text{cov}(X_i, S_i) & \text{var}(S_i)\end{pmatrix}.$$

We will first compute $\operatorname{cov}\big(X_i, Z_i^j\big)$ (for a fixed $j$) by conditionning on the value $IBD_j$, the number of alleles shared identical by descent by the index individual and the $j$-th sib. We have for $k = 0, 1, 2$,

$$E(X_i \mid IBD_j = k) = E\big(Z_i^j \mid IBD_j = k\big) = 0,$$

and

$$\operatorname{cov}\big(X_i, Z_i^j \mid IBD_j = k\big) = \begin{cases} \operatorname{cov}\big(X_i, Y_i^p + Y_i^m\big) & \text{if } k = 0 \\ \operatorname{cov}\big(X_i, X_i^p + Y_i^m\big) \text{ or } \operatorname{cov}\big(X_i, Y_i^p + X_i^m\big) & \text{if } k = 1 \\ \operatorname{cov}\big(X_i, X_i^p + X_i^m\big) & \text{if } k = 0 \end{cases}$$

the value of $\operatorname{cov}\big(X_i, Y_i^p + Y_i^m\big)$ is infinitesimal; we have $\operatorname{cov}\big(X_i, X_i^p + Y_i^m\big) = \operatorname{cov}\big(X_i, Y_i^p + X_i^m\big) \simeq p_i q_i$ and $\operatorname{cov}\big(X_i, X_i^p + X_i^m\big) = 2p_i q_i$. Using the law of total covariance and $\mathbb{P}(IBD_j = k) = \frac{1}{4}, \frac{1}{2}, \frac{1}{4}$ for $k = 0, 1, 2$, we get

$$\begin{aligned} \operatorname{cov}\big(X_i, Z_i^j\big) &= \sum_{k=0}^{2} \operatorname{cov}\big(X_i, Z_i^j \mid IBD = k\big)\mathbb{P}(IBD = k) \\ &\simeq \frac{1}{2} p_i q_i + \frac{1}{4} 2p_i q_i \\ &\simeq p_i q_i. \end{aligned}$$

From this we compute $\operatorname{var}(S_i)$ as a sum of $n$ terms equal to $2p_i q_i$ corresponding to $\operatorname{var}(X_i)$, $\operatorname{var}(Z_i^j)$, and $n(n-1)$ terms corresponding to $\operatorname{cov}(X_i, Z_i^j)$ or $\operatorname{cov}(Z_i^j, Z_i^{j'})$:

$$\operatorname{var}(S_i) = \frac{1}{n^2}\left(n \times 2p_i q_i + n(n-1) \times p_i q_i\right)$$

thus

$$\operatorname{var}(S_i) = \frac{n+1}{n} p_i q_i.$$

Similarly,

$$\operatorname{cov}(X_i, S_i) = \frac{1}{n}\left(\operatorname{var}(X_i) + \sum_j \operatorname{cov}\big(X_i, Z_i^j\big)\right)$$

which gives again

$$\operatorname{cov}(X_i, S_i) = \frac{n+1}{n} p_i q_i.$$

We have computed

$$\begin{pmatrix} \operatorname{var}(X_i) & \operatorname{cov}(X_i, S_i) \\ \operatorname{cov}(X_i, S_i) & \operatorname{var}(S_i) \end{pmatrix} = \frac{1}{n} p_i q_i \begin{pmatrix} 2n & n+1 \\ n+1 & n+1 \end{pmatrix}.$$

Using equation 91 we obtain

$$E\begin{pmatrix} \hat{\beta}_i \\ \hat{\beta}'_i \end{pmatrix} = \frac{1}{p_i q_i} \frac{n}{n-1} \begin{pmatrix} 1 & -1 \\ -1 & 2\frac{n}{n+1} \end{pmatrix} \begin{pmatrix} \operatorname{cov}(X_i, P) \\ \operatorname{cov}(S_i, P) \end{pmatrix},$$

and the expected value of the coefficient of $X_i$ in the regression of $P$ on $X_i$ and $S_i$ is

$$E\big(\hat{\beta}_i\big) = \frac{1}{p_i q_i} \frac{n}{n-1}\big(\operatorname{cov}(X_i, P) - \operatorname{cov}(S_i, P)\big).$$

or

$$E\left(\hat{\beta}_i\right) = \frac{1}{p_i q_i}\frac{n}{n-1}\operatorname{cov}(X_i - S_i, P). \tag{97}$$

We write $\operatorname{cov}(X_i, P) = 2\operatorname{cov}\left(X_i^p, P\right)$. We need to compute $\operatorname{cov}(S_i, P)$. We will again use the law of total covariance, with conditioning on $IBD_j$, to prove that $\operatorname{cov}(Z_i^j, P) = \operatorname{cov}\left(X_i^p, P\right) + \operatorname{cov}\left(Y_i^p, P\right)$, which will allow us to conclude the computation.

We first have

$$E\left(Z_i^j \mid IBD_j = k\right) = E(P \mid IBD_j = k) = 0,$$

and

$$\operatorname{cov}\left(Z_i^j, P \mid IBD_j = k\right) = \begin{cases} \operatorname{cov}\left(Y_i^p + Y_i^m, P\right) & \text{if } k = 0 \\ \operatorname{cov}\left(X_i^p + Y_i^m, P\right) \text{ or } \operatorname{cov}\left(Y_i^p + X_i^m, P\right) & \text{if } k = 1 \\ \operatorname{cov}\left(X_i^p + X_i^m, P\right) & \text{if } k = 0 \end{cases}$$

We have $\operatorname{cov}\left(Y^p, P\right) = \operatorname{cov}\left(Y_i^m, P\right)$ and $\operatorname{cov}\left(X_i^p, P\right) = \operatorname{cov}\left(X_i^m, P\right)$, so

$$\operatorname{cov}\left(Z_i^j, P \mid IBD_j = k\right) = \begin{cases} 2\operatorname{cov}\left(Y_i^p, P\right) & \text{if } k = 0 \\ \operatorname{cov}\left(X_i^p, P\right) + \operatorname{cov}\left(Y_i^p, P\right) & \text{if } k = 1 \\ 2\operatorname{cov}\left(X_i^p, P\right) & \text{if } k = 0 \end{cases}$$

and by the law of total covariance

$$\begin{aligned} \operatorname{cov}\left(P, Z_i^j\right) &= \sum_{k=0}^{2} \operatorname{cov}(P, Z_{1i} \mid IBD = k)\mathbb{P}(IBD = k) \\ &= \frac{1}{4} \times 2\operatorname{cov}\left(Y_i^p, P\right) + \frac{1}{2} \times \left(\operatorname{cov}\left(X_i^p, P\right) + \operatorname{cov}\left(Y_i^p, P\right)\right) + \frac{1}{4} \times 2\operatorname{cov}\left(X_i^p, P\right) \\ &= \operatorname{cov}\left(X_i^p, P\right) + \operatorname{cov}\left(Y_i^p, P\right), \end{aligned}$$

where we have used that $\operatorname{cov}\left(X_i^p, P\right) = \operatorname{cov}\left(X_i^m, P\right)$ and $\operatorname{cov}\left(Y_i^p, P\right) = \operatorname{cov}\left(Y_i^m, P\right)$.

We write

$$X_i - S_i = \frac{1}{n}\left((n-1)X_i - \sum_{j=1}^{n-1} Z_i^j\right),$$

thus

$$\begin{aligned} \operatorname{cov}(X_i - S_i, P) &= \frac{n-1}{n}\operatorname{cov}(X_i, P) - \frac{1}{n}\sum_{j=1}^{n-1}\operatorname{cov}\left(Z_i^j, P\right) \\ &= \frac{n-1}{n}\left(\operatorname{cov}(X_i, P) - \left(\operatorname{cov}\left(X_i^p, P\right) + \operatorname{cov}\left(Y_i^p, P\right)\right)\right) \\ &= \frac{n-1}{n}\left(\operatorname{cov}\left(X_i^p, P\right) - \operatorname{cov}\left(Y_i^p, P\right)\right) \end{aligned}$$

and, by equation 96,

$$\operatorname{cov}(X_i - S_i, P) = \frac{n-1}{n}\beta_i p_i q_i.$$

Using equation 97, we get that we have $E\left(\hat{\beta}_i\right) = \beta_i$, as announced.

##### 7.5 Sibling differences in phenotypes vs differences in genetic values

We consider pairs of siblings in a population at equilibrium. We denote by $A = G^p + G^m$ and $P = A + E$ the genetic value and the phenotype of the first sibling, and by $A_s = G_s^p + G_s^m$ and $P_s = A_s + E_s$ the corresponding quantities for the second sibling.

We are interested in the joint distribution of $A$ and $A_s$, of $P$ and $P_s$, and finally, in the joint distribution of $(A - A_s)$ and $(P - P_s)$. We will also consider the case where $A - A_s$ is substituted by $\hat{A} - \hat{A}_s$.

###### 7.5.1 Joint distribution of the genetic values

We have $\mathrm{var}(A) = \mathrm{var}(A_s) = a^2 = 2(1 + r_{\mathrm{ga}})g^2$ (equation 7) with $g^2 = \frac{g_0^2}{1 - r_{\mathrm{ga}}}$ (equation 33).

To fully specify the joint distribution of $A$ and $A_s$, we need $\mathrm{cov}(A, A_s)$. We write

$$\mathrm{cov}(A, A_s) = \mathrm{cov}\big(G^p, G_s^p\big) + \mathrm{cov}\big(G^p, G_s^m\big) + \mathrm{cov}\big(G^m, G_s^p\big) + \mathrm{cov}\big(G^m, G_s^m\big).$$

We have $\mathrm{cov}\big(G^p, G_s^m\big) = \mathrm{cov}\big(G^m, G_s^p\big) = r_{\mathrm{ga}} g^2$, as $r_{\mathrm{ga}}$ is the correlation between any two gametes emitted by the two mates. We just need $\mathrm{cov}\big(G^p, G_s^p\big)$ and $\mathrm{cov}\big(G^m, G_s^m\big)$, which are equal.

The distribution of $G^p$ conditional to the parental genetic value $A^p$ is Gaussian, with expected value $\frac{1}{2}A^p$ and variance $\frac{1}{2}(1 - r_{\mathrm{ga}})g^2$ (equation 12). Moreover, the covariance of $G^p$ and $G_s^p$ conditionnal to $A^p$ is 0. The law of total covariance implies

$$\mathrm{cov}\big(G^p, G_s^p\big) = \mathrm{cov}\left(\frac{1}{2}A^p, \frac{1}{2}A^p\right),$$

that is

$$\mathrm{cov}\big(G^p, G_s^p\big) = \frac{1}{2}(1 + r_{\mathrm{ga}})g^2. \tag{98}$$

This implies $\mathrm{cor}(G, G_s) = \frac{1}{2}(1 + r_{\mathrm{ga}})$.

Now turn back to $\mathrm{cov}(A, A_s)$: we have $\mathrm{cov}(A, A_s) = (1 + r_{\mathrm{ga}})g^2 + 2r_{\mathrm{ga}}g^2$, that is

$$\mathrm{cov}(A, A_s) = (1 + 3r_{\mathrm{ga}})g^2. \tag{99}$$

###### 7.5.2 Joint distribution of the environments

We have $\mathrm{var}(E) = \mathrm{var}(E_s) = e^2$. We need the value of $\mathrm{cov}(E, E_s)$.

The variance-covariance matrix of $(E^p, E^m, E)$ or $(E^p, E^m, E_s)$ is (equation 25)

$$e^2 \begin{pmatrix} 1 & r_{E^pE^m} & v \\ r_{E^pE^m} & 1 & v \\ v & v & 1 \end{pmatrix}$$

where $r_{E^pE^m} = \frac{r_{\mathrm{ho}}}{\sigma^2}(\rho a + e)^2$ (table 1, where it is denoted $r_{E_1E_2}$). This implies that the expected value of $E$ and $E_s$ conditional to $E^p, E^m$ is

$$E\big(E \mid E^p, E^m\big) = E\big(E_s \mid E^p, E^m\big) = \frac{v}{1 + r_{E^pE^m}}\big(E^p + E^m\big)$$

and variance $\left(1-2\frac{v^2}{1+r_{E^pE^m}}\right)e^2$. If we assume that the components of $E$ and $E_s$ that are not inherited from the parental environments $E^p$ and $E^m$ are independent, we have $\text{cov}\big(E,E_s \mid E^p,E^m\big)=0$.

The law of total covariance gives

$$\begin{aligned}
\text{cov}(E,E_s) &= E\big(\text{cov}\big(E,E_s \mid E^p,E^m\big)\big)+\text{cov}\big(E\big(E\mid E^p,E^m\big),E\big(E_s\mid E^p,E^m\big)\big)\\
&= \text{cov}\left(\frac{v}{1+r_{E^pE^m}}\big(E^p+E^m\big),\frac{v}{1+r_{E^pE^m}}\big(E^p+E^m\big)\right)\\
&= \frac{v^2}{(1+r_{E^pE^m})^2}\,\text{var}\big(E^p+E^m\big)\\
&= \frac{v^2}{(1+r_{E^pE^m})^2}\times 2(1+r_{E^pE^m})e^2
\end{aligned}$$

so

$$\text{cov}(E,E_s)=2\frac{v^2}{1+r_{E^pE^m}}e^2. \tag{100}$$

##### 7.5.3 Joint distribution of the phenotypes

We have naturally $\text{var}(P)=\text{var}(P_s)=\sigma^2$, and it is now easy to compute $\text{cov}(P,P_s)$. We have

$$\text{cov}(P,P_s)=\text{cov}(A,A_s)+\text{cov}(A,E_s)+\text{cov}(E,A_s)+\text{cov}(E,E_s).$$

By symmetry, we have $\text{cov}(A,E_s)=\text{cov}(E,A_s)=\rho ae$, or

$$\text{cor}(A,E_s)=\text{cor}(A_s,E)=\rho. \tag{101}$$

We have computed $\text{cov}(A,A_s)$ (equation 99) and $\text{cov}(E,E_s)$ (equation 100), so we can compute

$$\text{cov}(P,P_s)=(1+3r_{\text{ga}})g^2+2\rho ae+2\frac{v^2}{1+r_{E^pE^m}}e^2. \tag{102}$$

##### 7.5.4 Joint distribution of $(P-P_s)$ and $(A-A_s)$

Let us first compute $\text{cov}(P-P_s,A-A_s)$.

$$\begin{aligned}
\text{cov}(P-P_s,A-A_s) &= \text{cov}(P,A)-\text{cov}(P,A_s)-\text{cov}(P_s,A)+\text{cov}(P_s,A_s)\\
&= 2\,\text{cov}(P,A)-2\,\text{cov}(P,A_s).
\end{aligned}$$

The first term is

$$\begin{aligned}
\text{cov}(P,A) &= \text{var}(A)+\text{cov}(E,A)\\
&= a^2+\rho ae\\
&= 2(1+r_{\text{ga}})g^2+\rho ae.
\end{aligned}$$

The second term is (equation 99)

$$\text{cov}(P,A_s)=\text{cov}(A,A_s)+\text{cov}(E,A_s)\quad=(1+3r_{\text{ga}})g^2+\rho ae.$$

So we have

$$\operatorname{cov}(P - P_s, A - A_s) = 2(1 - r_{\text{ga}})g^2 = 2g_0^2. \tag{103}$$

We can also compute $\operatorname{var}(A - A_s)$:

$$\begin{aligned}\operatorname{var}(A - A_s) &= 2\operatorname{var}(A) - 2\operatorname{cov}(A, A_s) \\ &= 4(1 + r_{\text{ga}})g^2 - 2(1 + 3r_{\text{ga}})g^2\end{aligned}$$

and

$$\operatorname{var}(A - A_s) = 2(1 - r_{\text{ga}})g^2 = 2g_0^2. \tag{104}$$

And now, $\operatorname{var}(P - P_s)$:

$$\begin{aligned}\operatorname{var}(P - P_s) &= 2\operatorname{var}(P) - 2\operatorname{cov}(P, P_s) \\ &= 2\big(2(1 + r_{\text{ga}})g^2 + 2\rho ae + e^2\big) - 2\left((1 + 3r_{\text{ga}})g^2 + 2\rho ae + 2\frac{v^2}{1 + r_{E^pE^m}}e^2\right) \\ &= 2(1 - r_{\text{ga}})g^2 + 2\left(1 - 2\frac{v^2}{1 + r_{E^pE^m}}\right)e^2\end{aligned}$$

and

$$\operatorname{var}(P - P_s) = 2g_0^2 + 2\left(1 - 2\frac{v^2}{1 + r_{E^pE^m}}\right)e^2. \tag{105}$$

Thus

$$\operatorname{cor}(P - P_s, A - A_s)^2 = \frac{2g_0^2}{2g_0^2 + 2\left(1 - 2\frac{v^2}{1 + r_{E^pE^m}}\right)e^2}. \tag{106}$$

Remark that in the absence of VCT, that is if $v = 0$, writing $h_0^2 = \frac{2g_0^2}{2g_0^2 + e^2}$, we get

$$\operatorname{cor}(P - P_s, A - A_s)^2 = \frac{2h_0^2}{2 - h_0^2},$$

which gives a one-to-one correspondance between $h_0^2$ and $\operatorname{cor}(P - P_s, A - A_s)^2$.

##### 7.5.5 Using the Polygenic Scores

We are going to show briefly in this section than when $A - A_s$ is replaced by $\hat{A} - \hat{A}_s$, the correlation in equation 106 has to be corrected by a mutliplicative factor, namely:

$$\operatorname{cor}\left(P - P_s, \hat{A} - \hat{A}_s\right)^2 = \operatorname{cor}(P - P_s, A - A_s)^2 \frac{\gamma^2}{\gamma^2 + \frac{N}{M}\frac{\sigma^2}{g_0^2}} \tag{107}$$

We fill first show that

$$\operatorname{cov}\left(P - P_s, \hat{A} - \hat{A}_s\right) = \gamma \operatorname{cov}(P - P_s, A - A_s) \tag{108}$$

We write this as

$$\begin{aligned}\operatorname{cov}\left(P - P_s, \hat{A} - \hat{A}_s\right) &= \operatorname{cov}\left(P, \hat{A}\right) + \operatorname{cov}\left(P_s, \hat{A}_s\right) - \operatorname{cov}\left(P, \hat{A}_s\right) - \operatorname{cov}\left(P_s, \hat{A}\right) \\ &= 2\operatorname{cov}\left(P, \hat{A}\right) - 2\operatorname{cov}\left(P, \hat{A}_s\right)\end{aligned}$$

as the two sibs play symmetric role.

We have, from equations 77 and 13,

$$\operatorname{cov}\left(P, \hat{A}\right) = \operatorname{cov}(P_s, A_s) = \gamma a(a + \rho e) = \gamma \operatorname{cov}(P, A).$$

We need to show that $\operatorname{cov}\left(P, \hat{A}_s\right) = \gamma \operatorname{cov}(P, A_s)$. To verify this, we write $\operatorname{cov}\left(P, \hat{A}_s\right)$ as

$$\operatorname{cov}\left(P, \hat{A}_s\right) = \operatorname{cov}\left(A, \hat{A}_s\right) + \operatorname{cov}\left(E, \hat{A}_s\right).$$

Let us show that $\operatorname{cov}\left(A, \hat{A}_s\right) = \gamma \operatorname{cov}(A, A_s)$. We have

$$\begin{aligned}
\operatorname{cov}\left(A, \hat{A}_s\right) &= \operatorname{cor}\left(A, \hat{A}_s\right)\sqrt{\operatorname{var}(A)\operatorname{var}\left(\hat{A}_s\right)} \\
&= \operatorname{cor}(A, A_s)\operatorname{cor}\left(A_s, \hat{A}_s\right) a\sqrt{\gamma^2 a^2 + \frac{N}{M}\sigma^2} && \text{(from equation 71)} \\
&= \operatorname{cor}(A, A_s) a^2 \gamma && \text{(from equation 73)} \\
&= \gamma \operatorname{cov}(A, A_s),
\end{aligned}$$

as $\operatorname{var}(A) = \operatorname{var}(A_s) = a^2$.

Next let us show that $\operatorname{cov}\left(E, \hat{A}_s\right) = \gamma \operatorname{cov}(E, A_s)$. We have

$$\begin{aligned}
\operatorname{cov}\left(E, \hat{A}_s\right) &= \operatorname{cor}\left(E, \hat{A}_s\right)\sqrt{\operatorname{var}(E)\operatorname{var}\left(\hat{A}_s\right)} \\
&= \operatorname{cor}(E, A_s)\operatorname{cor}\left(A_s, \hat{A}_s\right) e\sqrt{\gamma^2 a^2 + \frac{N}{M}\sigma^2} && \text{(from equation 71)} \\
&= \operatorname{cor}(E, A_s)\gamma a e && \text{(from equation 73)} \\
&= \gamma \operatorname{cov}(E, A_s).
\end{aligned}$$

Putting these two results together, we get $\operatorname{cov}\left(P, \hat{A}_s\right) = \gamma \operatorname{cov}(P, A_s)$. Finally we have

$$\begin{aligned}
\operatorname{cov}\left(P - P_s, \hat{A} - \hat{A}_s\right) &= 2\operatorname{cov}\left(P, \hat{A}\right) - 2\operatorname{cov}\left(P, \hat{A}_s\right) \\
&= 2\gamma \operatorname{cov}(P, A) - 2\gamma \operatorname{cov}(P, A_s)
\end{aligned}$$

and

$$\operatorname{cov}\left(P - P_s, \hat{A} - \hat{A}_s\right) = \gamma \operatorname{cov}(P - P_s, A - A_s). \tag{109}$$

To conclude, we need to compute $\operatorname{var}\left(\hat{A} - \hat{A}_s\right)$. We have

$$\begin{aligned}
\operatorname{var}\left(\hat{A} - \hat{A}_s\right) &= \operatorname{var}\left(\hat{A}\right) - 2\operatorname{cov}\left(\hat{A}, \hat{A}_s\right) + \operatorname{var}\left(\hat{A}_s\right) \\
&= 2\operatorname{var}\left(\hat{A}\right) - 2\operatorname{cov}\left(\hat{A}, \hat{A}_s\right) \\
&= 2\left(\gamma^2 a^2 + \frac{N}{M}\sigma^2\right) - 2\operatorname{cov}\left(\hat{A}, \hat{A}_s\right)
\end{aligned}$$

(from equation 71). Write

$$\begin{aligned}
\operatorname{cor}\left(\hat{A}, \hat{A}_s\right) &= \operatorname{cor}(A, A_s)\operatorname{cor}\left(A, \hat{A}\right)\operatorname{cor}\left(A_s, \hat{A}_s\right) \\
&= \operatorname{cor}(A, A_s)\frac{\gamma^2 a^2}{\gamma^2 a^2 + \frac{N}{M}\sigma^2}
\end{aligned}$$

from equation 73. Thus

$$\begin{aligned}
\operatorname{cov}\left(\hat{A},\hat{A}_s\right) &= \operatorname{cor}(\left(\hat{A},\hat{A}_s\right)\operatorname{var}(A) \\
&= \operatorname{cor}(A,A_s)\frac{\gamma^2 a^2}{\gamma^2 a^2+\frac{N}{M}\sigma^2}\left(\gamma^2 a^2+\frac{N}{M}\sigma^2\right) && \text{(from equation 71)} \\
&= \gamma^2 a^2 \operatorname{cor}(A,A_s) \\
&= \gamma^2 \operatorname{cov}(A,A_s) \\
&= \gamma^2(1+3r_{\text{ga}})g^2. && \text{(from equation 99)}
\end{aligned}$$

So we have

$$\begin{aligned}
\operatorname{var}\left(\hat{A}-\hat{A}_s\right) &= 2\left(\gamma^2 a^2+\frac{N}{M}\sigma^2\right)-2\operatorname{cov}\left(\hat{A},\hat{A}_s\right) \\
&= 2\left(\gamma^2 a^2+\frac{N}{M}\sigma^2\right)-2\gamma^2(1+3r_{\text{ga}})g^2 \\
&= 2\gamma^2\left(a^2-(1+3r_{\text{ga}})g^2\right)+2\frac{N}{M}\sigma^2
\end{aligned}$$

But now we have $a^2=\frac{1+r_{\text{ga}}}{1-r_{\text{ga}}}2g_0^2$ (equation 40) and $g^2=\frac{g_0^2}{1-r_{\text{ga}}}$ (equation 39) . Hence

$$\begin{aligned}
\operatorname{var}\left(\hat{A}-\hat{A}_s\right) &= 2\gamma^2\frac{1}{1-r_{\text{ga}}}g_0^2\left(2(1+r_{\text{ga}})-(1+3r_{\text{ga}})\right)+2\frac{N}{M}\sigma^2 \\
&= 2\gamma^2 g_0^2+2\frac{N}{M}\sigma^2.
\end{aligned}$$

As we have $\operatorname{var}(A-A_s)=2g_0^2$ (equation 104), we deduce that

$$\operatorname{var}\left(\hat{A}-\hat{A}_s\right)=\operatorname{var}(A-A_s)\left(\gamma^2+\frac{N}{M}\frac{\sigma^2}{g_0^2}\right). \tag{110}$$

Finally, using equations 109 and 110 together, we get

$$\begin{aligned}
\operatorname{cor}\left(P-P_s,\hat{A}-\hat{A}_s\right)^2 &= \frac{\operatorname{cov}\left(P-P_s,\hat{A}-\hat{A}_s\right)^2}{\operatorname{var}(P-P_s)\operatorname{var}\left(\hat{A}-\hat{A}_s\right)} \\
&= \frac{\gamma^2\operatorname{cov}(P-P_s,A-A_s)^2}{\operatorname{var}(P-P_s)\operatorname{var}(A-A_s)\left(\gamma^2+\frac{N}{M}\frac{\sigma^2}{g_0^2}\right)} \\
&= \operatorname{cor}(P-P_s,A-A_s)^2\frac{\gamma^2}{\gamma^2+\frac{N}{M}\frac{\sigma^2}{g_0^2}}.
\end{aligned}$$

#### 7.6 Untransmitted alleles

When parental genotypes are available, one might be interested in investigating the 'Nature of Nurture' hypothesis of parental (untransmitted) genotypes having indirect genetic effects on the phenotype of the offspring. Hereafter, we first compute the correlation between the phenotype and the so-called "untransmitted genetic value" (or its estimate by a PGS) ; then we show that the regression on untransmitted alleles provides a way to obtain unbiased estimates of the allelic effect.

##### 7.6.1 Correlation between the phenotype and the untransmitted genetic value

If the untransmitted gametes in a trio are labeled as $H^p$ and $H^m$, then let $A_u = H^p + H^m$ be the untransmitted genetic value in the trio. We have

$$\operatorname{cov}(A_u, P) = \operatorname{cov}\left(H^p + H^m, P\right) = \operatorname{cov}\left(H^p + H^m, G^p + G^m + E\right);$$

by symmetry of the maternal and paternal indexes $m$ and $p$, this will be:

$$\operatorname{cov}\left(A_u, P\right) = 2\operatorname{cov}\left(H^p, G^p\right) + 2\operatorname{cov}\left(H^p, G^m\right) + 2\operatorname{cov}\left(H^p, E\right).$$

Both $\operatorname{cov}\left(H^p, G^p\right)$ and $\operatorname{cov}\left(H^p, G^m\right)$ are simply equal to $r_{\text{ga}} g^2$ if the population is at equilibrium. As $H^p$ and $H^m$ play symmetric roles, we have $\operatorname{cov}\left(H^p, E\right) = \frac{1}{2}\operatorname{cov}(A_u, E)$, and this is in turn equal to $\frac{1}{2}\operatorname{cov}(A, E) = \frac{1}{2}\rho a e$, given that $A_u$ has all the properties of $A$. Hence we finally have:

$$\operatorname{cov}\left(H^p + H^m, P\right) = \operatorname{cov}(A_u, P) = 4 r_{\text{ga}} g^2 + \rho a e.$$

Hence, the correlation between $A_u$ and $P$ can be calculated, and will be:

$$\operatorname{cor}(A_u, P) = \frac{4 r_{\text{ga}} g^2 + \rho a e}{a\sigma},$$

which becomes, using equation 31 and 34:

$$\operatorname{cor}(A_u, P) = \frac{r_{A_1 A_2} a}{\sigma} + \frac{\rho e}{\sigma}.$$

There are in this correlation two interpretable components, coming respectively from AM and VCT.

###### Using a polygenic score

If the same correlation were to be calculated using a polygenic score calculated in the untransmitted alleles in the trio, $\hat{A}_u$, then as $P$ and $\hat{A}_u$ are conditionally independant given $A_u$, we can use equation 76 to get:

$$\operatorname{cor}(\hat{A}_u, P) = \sqrt{\frac{h^2_{SNP}}{h^2_{SNP} + \frac{N}{M}\left(1 - r_{\text{ho}} h^2_{SNP}\right)^2}} \cdot \left(\frac{r_{A_1 A_2} a}{\sigma} + \frac{\rho e}{\sigma}\right).$$

##### 7.6.2 Regression on untransmitted alleles

We can also consider the association statistic coming from an 'indirect effect' GWAS where untransmitted genotypes are regressed against offspring phenotype. This leads to a similar calculation as in section 6.2, but now we would be interested in calculating $\text{cov}(X_{iu}, P)$ where $X_{iu} = X_{iu}^p + X_{iu}^m$ is the untransmitted alleles for variant $i$. This covariance is again composed of three parts, as described in sections 6.2.1, 6.2.2, and 6.2.3.

For the equivalent calculation of 6.2.1, we are now interested in $\text{cov}(X_{iu}^p, G^p)$ which will be simply equal to $r_{\text{ga}}\,\text{cov}(X_{iu}^p, G^p)$ given that $X_{iu}^p$ and $G^p$ are conditionally independent given $H^p$, the complete untransmitted paternal genetic value, and hence

$$\text{cor}(X_{iu}^p, G^p) = \text{cor}(X_{iu}^p, H^p)\,\text{cor}(G^p, H^p),$$

and $\text{cor}(X_{iu}^p, H^p) = \text{cor}(X_i^p, G^p)$ by symmetry and $\text{cor}(G^p, H^p) = r_{\text{ga}}$.

For the calculation in 6.2.2, we are now interested in $\text{cov}(X_{iu}^p, G^m)$, and we have

$$\text{cov}(X_{iu}^p, G^m) = \text{cov}(X_i^p, G^m),$$

taking advantage of the randomness of transmission.

The same holds for equivalent calculation of 6.2.3, we now need to calculate $\text{cov}(X_{iu}^p, E)$ which is again equal to $\text{cov}(X_i^p, E)$ again by randomness of transmission.

The upshot is that we can follow the exact same logic as section 6.2, until we reach equation 63 which becomes:

$$\text{cov}(X_i^p, P) = \left(r_{\text{ga}} + r_{\text{ga}} + \rho\frac{e}{a}(1 + r_{\text{ga}})\right)\text{cov}(X_i^p, G^p)$$

where the only difference is that the first term in the brackets has changed from 1 to $r_{\text{ga}}$. This leads, using the same logic as 6.2.4, to the expected regression coefficient from a GWAS on untransmitted alleles being:

$$\begin{aligned} E\left(\hat{\beta}_{iu}\right) &= 2\left(2r_{\text{ga}} + \rho\frac{e}{a}(1 + r_{\text{ga}})\right)\beta_i p_i q_i . \frac{1}{2p_i q_i (1 - r_{\text{ga}})} \\ &= \frac{1 + r_{\text{ga}}}{1 - r_{\text{ga}}}\left(\frac{2r_{\text{ga}}}{1 + r_{\text{ga}}} + \rho\frac{e}{a}\right)\beta_i \\ &= \frac{1 + r_{\text{ga}}}{1 - r_{\text{ga}}}\left(r_{A_1 A_2} + \rho\frac{e}{a}\right)\beta_i \\ &= (\gamma - 1)\beta_i, \end{aligned}$$

using again 31 and 66.

Hence, for a variant that is causal, the difference between association statistics between on transmitted and untransmitted alleles would be:

$$E\left(\hat{\beta}_i - \hat{\beta}_{iu}\right) = \gamma\beta_i - (\gamma - 1)\beta_i = \beta_i,$$

providing another route to an unbiased estimate of $\beta_i$ under our model of AM with VCT.

### A Technical lemma

If $A, B, X$ are three random variables, the partial correlation $r_{AB\cdot X}$ is the correlation of $A$ and $B$ conditional to $X$.

**Lemma 1 (Partial correlation)** *Let $(A, B, X)$ be a Gaussian vector. If $A$ and $B$ are independent conditional to $X$, or equivalently $r_{AB\cdot X} = 0$, then $r_{AB} = r_{AX} r_{BX}$.*

**Proof** Assume without lost of generality $\mathrm{var}(A) = \mathrm{var}(B) = \mathrm{var}(X) = 1$. The variance-covariance matrix of $(A, B, X)$ is

$$\mathrm{var}(A, B, X) = \begin{pmatrix} 1 & r_{AB} & r_{AX} \\ r_{AB} & 1 & r_{BX} \\ r_{AX} & r_{BX} & 1 \end{pmatrix}.$$

Standard Gaussian theory allows to compute $\mathrm{var}(A, B \mid X)$:

$$\mathrm{var}(A, B \mid X) = \begin{pmatrix} 1 & r_{AB} \\ r_{AB} & 1 \end{pmatrix} - \begin{pmatrix} r_{AX} \\ r_{BX} \end{pmatrix} \begin{pmatrix} r_{AX} & r_{BX} \end{pmatrix} = \begin{pmatrix} 1 - r_{AX}^2 & r_{AB} - r_{AX} r_{BX} \\ r_{AB} - r_{AX} r_{BX} & 1 - r_{BX}^2 \end{pmatrix},$$

implying $r_{AB\cdot X} = r_{AB} - r_{AX} r_{BX}$. □

**Lemma 2** *Let $(X_1, X_2, Y_1, Y_2)$ be a gaussian vector. Note $X = (X_1, X_2)$ and $Y = (Y_1, Y_2)$. Assume*

$$\mathrm{var}(Y) = \begin{pmatrix} 1 & \alpha \\ \alpha & 1 \end{pmatrix}$$

*and*

$$\mathrm{cov}(X, Y) = \begin{pmatrix} u & v \\ w & w \end{pmatrix}.$$

*If $X_1, X_2$ are independent conditionally to $Y$, or $\mathrm{cov}(X_1, X_2 \mid Y) = 0$, then*

$$\mathrm{cov}(X_1, X_2) = \frac{u + v}{1 + \alpha} w.$$

**Proof** The variance-covariance matrix of $X$ conditional to $Y$ is

$$\mathrm{var}(X) - \mathrm{cov}(X, Y)\,\mathrm{var}(Y)^{-1}\,\mathrm{cov}(Y, X) = \frac{1}{1 - \alpha^2} \begin{pmatrix} (1-\alpha)2uv + (u - v)^2 & (1-\alpha)(u + v)w \\ (1-\alpha)(u + v)w & (1-\alpha)2w^2 \end{pmatrix}$$

from which the result is obtained readily. □

**Lemma 3** *Let $A_n$ and $X_n$ be sequences of random variables. For all $n$, $A_n$ is binary, that is, it takes only two different values, and $X_n$ is Gaussian. Both $A_n$ and $X_n$ have expected value equal to $0$ and variance equal to $1$. Aussume that for all $n$*

$$\mathrm{cov}(A_n, X_n) = \frac{1}{n} d.$$

*Then, in probability,*

$$nE(A_n \mid X_n) \underset{n\to\infty}{\longrightarrow} dX_n.$$

*This means in practice that for $n$ large enough, $E(A_n \mid X_n) \simeq \mathrm{cov}(A_n, X_n) X_n$.*

**Proof** As $A_n$ takes only two values, $E(X_n \mid A_n)$ can be written as a linear function of $A_n$. We let $E(X_n \mid A_n) = a + bX_n$. As $E(X_n) = E(a + bA_n) = a = 0$, we have simply $E(X_n \mid A_n) = bA_n$. Moreover $E(X_n A_n \mid A_n) = bA_n^2$, and taking expected values on both sides of this equation leads to $E(X_n A_n) = b\operatorname{var}(A_n) = b$, thus $b = \operatorname{cov}(X_n, A_n) = \frac{1}{n}d$. We have proved that

$$E(X_n \mid A_n) = \frac{d}{n} A_n.$$

For $n$ large, as $\operatorname{cov}(X_n, A_n)$ becomes infinitesimal, the distribution of $X_n$ conditional to $A_n$ will be approximately Gaussian. From $\operatorname{var}(X_n) = E(\operatorname{var}(X_n \mid A_n)) + \operatorname{var}(E(X_n \mid A_n))$, we get $\operatorname{var}(X_n \mid A_n) = 1 - \frac{d^2}{n^2}$. Now

$$\begin{aligned}
E(A_n \mid X_n = x) &= \sum_a aP(A_n = a \mid X_n = x) \\
&= \sum_a a \frac{P(X_n = x \mid A_n = a)}{\mathbb{P}(X_n = x)} \mathbb{P}(A_n = a) \\
&= \sum_a a\mathbb{P}(A = a) \left(1 - \frac{d^2}{n^2}\right)^{-\frac{1}{2}} \exp\left(-\frac{1}{2}\left(\frac{\left(x - \frac{d}{n}a\right)^2}{1 - \frac{d^2}{n^2}} - x^2\right)\right).
\end{aligned}$$

Let $f_n(x) = nE(A_n \mid X_n = x)$. We have

$$\begin{aligned}
f_n(0) &= n\sum_a a\mathbb{P}(A_n = a)\left(1 - \frac{d^2}{n^2}\right)^{-\frac{1}{2}} \exp\left(-\frac{1}{2}\frac{\frac{d^2}{n^2}a^2}{1 - \frac{d^2}{n^2}}\right) \\
&= \sum_a a\mathbb{P}(A_n = a) n \left(1 + \frac{1}{2}\frac{d^2}{n^2} + O\left(\frac{1}{n^4}\right)\right)\left(1 - \frac{1}{2}\frac{d^2 a^2}{n^2} + O\left(\frac{1}{n^4}\right)\right) \\
&= \sum_a a\mathbb{P}(A_n = a) n \left(1 + \frac{1}{2}d^2\left(1 - a^2\right)\frac{1}{n^2} + O\left(\frac{1}{n^4}\right)\right) \\
&= \sum_a a\mathbb{P}(A_n = a)\left(n + \frac{1}{2}d^2\left(1 - a^2\right)\frac{1}{n} + O\left(\frac{1}{n^3}\right)\right) \\
&= \left(\sum_a a\mathbb{P}(A_n = a)\right) n + \frac{1}{n}\sum_a a\mathbb{P}(A_n = a)\left(\frac{1}{2}d^2\left(1 - a^2\right) + O\left(\frac{1}{n^2}\right)\right) \\
&= \frac{1}{n}\sum_a a\mathbb{P}(A_n = a)\left(\frac{1}{2}d^2\left(1 - a^2\right) + O\left(\frac{1}{n^2}\right)\right)
\end{aligned}$$

as $\sum_a a\mathbb{P}(A_n = a) = E(A_n) = 0$. We conclude that $f_n(0) \to 0$ when $n \to \infty$. Now we have

$$\begin{aligned}
f_n'(x) &= n\sum_a a\mathbb{P}(A_n = a)\left(1 - \frac{d^2}{n^2}\right)^{-\frac{1}{2}} \times \exp\left(-\frac{1}{2}\left(\frac{\left(x - \frac{d}{n}a\right)^2}{1 - \frac{d^2}{n^2}} - x^2\right)\right) \times \left(x - \frac{x - \frac{d}{n}a}{1 - \frac{d^2}{n^2}}\right) \\
&= \sum_a a\mathbb{P}(A_n = a)\left(1 - \frac{d^2}{n^2}\right)^{-\frac{1}{2}} \times \exp\left(-\frac{1}{2}\left(\frac{\left(x - \frac{d}{n}a\right)^2}{1 - \frac{d^2}{n^2}} - x^2\right)\right) \times \left(-\frac{\frac{d^2}{n}x}{1 - \frac{d^2}{n^2}} + \frac{da}{1 - \frac{d^2}{n^2}}\right)
\end{aligned}$$

and thus

$$f_n'(x) \underset{n\to\infty}{\longrightarrow} \sum_a aP(A_n = a) \times da = d\sum_a a^2 P(A_n = a) = dE(A_n^2) = d.$$

Finally, write $f_n(x) = f_n(0) + f_n'(t_x \cdot x)x$ with $t_x \in [0,1]$. We have

$$\sqrt{n}E(A_n \mid X_n) = f_n(0) + f_n'(t_{X_n} X_n)X_n \longrightarrow dX_n.$$

□

**Corollary 1** *Let $A_n$ and $X_n$ be as in lemma 3, and let $Y_n$ be a Gaussian variable, with $E(Y_n) = 0$, $\mathrm{var}(Y_n) = 0$, and $\mathrm{cor}(X_n, Y_n) = r$. Assume that $(X_n, Y_n)$ is bivariate Gaussian, and that $A_n$ independent of $Y_n$ conditionally to $X_n$. Then*

$$n\,\mathrm{cov}(A_n \mid Y_n) \underset{n\to\infty}{\longrightarrow} rd.$$

*In practice, for n large enough, $\mathrm{cov}(A_n, Y_n) \simeq r\,\mathrm{cov}(A_n, X_n)$.*

**Proof** We have $\mathrm{cov}(A_n, Y_n \mid X_n) = 0$ and $E(Y_n \mid X_n) = rX_n$. Lemma 3 tells that $nE(A_n \mid X_n) \to dX_n$ for $n \to \infty$. We get

$$\begin{aligned} n\,\mathrm{cov}(A_n, Y_n) &= nE(\mathrm{cov}(A_n, Y_n \mid X_n)) + \mathrm{cov}(nE(A_n \mid X_n), E(Y_n \mid X_n)) \\ &= \mathrm{cov}(nE(A_n \mid X_n), rX_n) \\ &= rE(nE(A_n \mid X_n)X_n) \to rE(dX_n^2) = rd. \end{aligned}$$

□

The following corollary establishes a result used in section 5. The notations used are those used along the whole text, and mostly defined in section 1.

**Corollary 2** *Let $g_i^p$, $G^p$, $g_j^m$, and $G^m$, be centered random variables, with $g_i^p$ and $g_i^m$ binary variables and $G^p$ and $G^m$ Gaussian variables. Assume that $\mathrm{var}(g_i^p) = \frac{1}{N}g_0^2\kappa_i^2$, $\mathrm{var}(G^p) = Ng_0^2\overline{\kappa}$, $\mathrm{cov}(g_i^p, G^p) = \frac{1}{N}g_0^2\overline{\kappa}_i$, and that $\mathrm{var}(g_j^m) = \frac{1}{N}g_0^2\kappa_j^2$, $\mathrm{var}(G^m) = Ng_0^2\overline{\kappa}$, $\mathrm{cov}(g_j^m, G^m) = \frac{1}{N}g_0^2\overline{\kappa}_j$. Assume that $\mathrm{cov}(G^p, G^m) = r_{ga}$ and that $g_i^p$, $g_j^m$ are independent conditionally to $G^m$. Then*

$$\mathrm{cov}(g_i^p, g_j^m) = \frac{1}{N}g_0^2 r_{ga}\frac{\overline{\kappa}_i\overline{\kappa}_j}{\overline{\kappa}}.$$

**Proof** Let $A = \frac{\sqrt{N}}{g_0\kappa_i}g_i^p$, so that $\mathrm{var}(A) = 1$. Let $X = \frac{1}{\sqrt{Ng_0^2\overline{\kappa}}}G^p$ so that $\mathrm{var}(X) = 1$. We have $\mathrm{cov}(A, X) = \frac{1}{g_0^2\kappa_i\sqrt{\overline{\kappa}}}\mathrm{cov}(g_i^p, G^p) = \frac{1}{N}\frac{\overline{\kappa}_i}{\kappa_i\sqrt{\overline{\kappa}}}$.

Let $Y = \frac{1}{\sqrt{Ng_0^2\overline{\kappa}}}G^m$. We have $\mathrm{cor}(X, Y) = r_{\mathrm{ga}}$. Applying corollary 1, for $N$ large enough we have $\mathrm{cov}(A, Y) \simeq r_{\mathrm{ga}}\,\mathrm{cov}(A, X) = \frac{1}{N}r_{\mathrm{ga}}\frac{\overline{\kappa}_i}{\kappa_i\sqrt{\overline{\kappa}}}$.

Now let $B = \frac{\sqrt{N}}{g_0\kappa_j}g_j^m$. We have $\mathrm{cov}(B, Y) = \frac{1}{N}\frac{\overline{\kappa}_j}{\kappa_j\sqrt{\overline{\kappa}}}$. Let's compute $\mathrm{cov}(A, B)$ using the law of total covariance:

$$\begin{aligned} \mathrm{cov}(A, B) &= E(\mathrm{cov}(A, B \mid Y)) + \mathrm{cov}(E(A \mid Y), E(B \mid Y)) \\ &= \mathrm{cov}(E(A \mid Y), E(B \mid Y)) \end{aligned}$$

as $A$ and $B$ are independent conditionally to $Y$. Using lemma 3, we have $E(A \mid Y) \simeq \operatorname{cov}(A, Y) Y = \frac{1}{N} r_{\text{ga}} \frac{\overline{\kappa}_i}{\kappa_i \sqrt{\overline{\kappa}}} Y$ and $E(B \mid Y) \simeq \operatorname{cov}(B, Y) Y = \frac{1}{N} \frac{\overline{\kappa}_j}{\kappa_j \sqrt{\overline{\kappa}}}$. It comes

$$\begin{aligned}\operatorname{cov}(A, B) &= \frac{1}{N^2} r_{\text{ga}} \frac{\overline{\kappa}_i \overline{\kappa}_j}{\kappa_i \kappa_j \overline{\kappa}} \operatorname{cov}(Y, Y) \\ &= \frac{1}{N^2} r_{\text{ga}} \frac{\overline{\kappa}_i \overline{\kappa}_j}{\kappa_i \kappa_j \overline{\kappa}}\end{aligned}$$

Finally, as $\operatorname{cov}(A, B) = \frac{N}{g_0^2 \kappa_i \kappa_j} \operatorname{cov}\left(g_i^p, g_j^m\right)$, we have

$$\operatorname{cov}\left(g_i^p, g_j^m\right) = \frac{1}{N} g_0^2 r_{\text{ga}} \frac{\overline{\kappa}_i \overline{\kappa}_j}{\overline{\kappa}}.$$

□

### B Computing the evolution of a population

Here we expose briefly how to use the results of sections 3 and 4 to compute the values of all parameters at a given generation $t$, assuming unlinked loci all contributing equally to the phenotype.

We consider a population described by the parameters $g_0$, $e$, $r_{\text{ho}}$ and $v$ ; as mentionnend at the beginning of section 4 these are the natural parameters, from which all other can be derived.

We assume that $r_{\text{ho}}$ and $v$ were both null before $t = 0$, so that we have $r_{\text{ga}}(0) = \rho(0) = \overline{\kappa}(0) = 0$, and $g(0) = g_0$. We show how to compute all parameters at time $t+1$, given their values at time $t$. At each $t$ we let $a(t)^2 = 2(1 + r_{\text{ga}}(t)) g(t)^2$ (equation 7) and $\sigma^2(t) = a(t)^2 + 2\rho(t) a(t) e + e^2$.

At time $t = 0$ we have $\kappa_{ii} = 1$ and $\kappa_{ij} = 0$ for $i \neq j$, thus $\overline{\kappa}(0) = \frac{1}{N}$. In the case of unlinked loci, the equation of evolution of the $\kappa_{ij}$ (equation 36) specializes to

$$\kappa_{ij}(t+1) = \frac{1}{2} \kappa_{ij}(t) + \frac{1}{2} r_{\text{ga}}(t) \overline{\kappa}(t)$$

for $i \neq j$ ; assumming all loci have equal variance we have $\kappa_{ii} = \kappa_i^2 = 1$. Using $N^2 \overline{\kappa}(t) = N + \sum_{i \neq j} \kappa_{ij}(t)$, we obtain

$$\overline{\kappa}(t+1) = \frac{1}{2N} + \frac{1}{2} \overline{\kappa}(t) + \frac{1}{2}\left(1 - \frac{1}{N}\right) r_{\text{ga}}(t) \overline{\kappa}(t).$$

As $\overline{\kappa}(t)$ is of order of magnitude $\frac{1}{N}$, one can prefer to handle $N\overline{\kappa}(t)$, which verifies $N\overline{\kappa}(0) = 1$ and

$$N\overline{\kappa}(t+1) = \frac{1}{2} + \frac{1}{2} N\overline{\kappa}(t) + \frac{1}{2}\left(1 - \frac{1}{N}\right) r_{\text{ga}}(t) N\overline{\kappa}(t).$$

Then we have (equation 6)

$$g(t+1)^2 = N\overline{\kappa}(t+1) g_0^2,$$

and (equations 23 and 16),

$$r_{\text{ga}}(t+1) = \frac{1}{2}\frac{r_{\text{ho}}}{\sigma^2(t)}(a(t)+\rho(t)e)^2\frac{g^2(t)}{g^2(t+1)}(1+r_{\text{ga}}(t)),$$

and (equation 29),

$$\rho(t+1) = \nu\frac{a(t)}{a(t+1)}\frac{\rho(t)+\frac{r_{\text{ho}}}{\sigma^2(t)}\big(\rho(t)a(t)+e\big)\big(\rho(t)e+a(t)\big)}{1+\frac{r_{\text{ho}}}{\sigma^2(t)}\big(\rho(t)a(t)+e\big)^2}.$$
